# Top-down inputs are controlled by somatostatin-expressing interneurons during associative learning

**DOI:** 10.1101/2024.12.18.629098

**Authors:** Florian Studer, Tommaso Zeppillo, Tania Rinaldi Barkat

## Abstract

Associative learning links sensory signals to their behavioral meaning by combining bottom-up inputs with top-down contextual information, enabling decision-making based on expected outcomes. It triggers plastic changes of neuronal responses in both excitatory and inhibitory cell populations. However, the role of inhibition in shaping this plasticity remains debated. Here we used chronic extracellular electrophysiology and optogenetic manipulation of inhibitory neurons and top-down inputs in mice learning an auditory discrimination task. We found that learning enhances stimulus selectivity in a subset of neurons in the primary auditory cortex. Interestingly, somatostatin-expressing (SST) interneurons decrease their response to the rewarded cue and bidirectionally regulate associative learning. More specifically, an increased activity of SST neurons impairs learning by altering bottom-up signaling, whereas the reduction of their activity facilitates learning by gating top-down inputs from the orbitofrontal cortex. These findings demonstrate that inhibition plays a critical role in gating top-down inputs to primary sensory cortices involved in associative learning.

## Introduction

Associative learning links sensory cues to their behavioral relevance and plays a crucial role in flexible behavioral adaptation. It enables the proper interpretation of our environment by comparing bottom-up sensory signals with top-down information related to the context and the valence of an action’s outcome ^1,2^. A substantial body of evidence suggests that learning shapes cortical circuits, reinforcing the neuronal networks involved in processing bottom-up signals ^3–6^. Improved behavioral performance – often regarded as the hallmark of learning - is suggested to be associated with enhanced neuronal discriminability for the learned stimulus ^7,8^.

Recent studies have shown that this increased discriminability is driven by synaptic plasticity in distal tuft dendrites of excitatory neurons within primary sensory cortices, mediated by inputs from higher-order brain areas ^9–12^. Inhibitory neurons have also been shown to be involved in experience-induced plasticity ^13^. Somatostatin-expressing (SST+) and NDNF-expressing interneurons exhibit changes in the strength of their responses to learned stimuli ^14,15^. Vaso-intestinal peptide expressing interneurons (VIP+) tend to show stronger responses to the outcomes of behavioral actions after learning ^16^. Additionally, inhibition is suggested to take part in associative learning not only in the cortex ^17–20^ but also in subcortical regions essential for aversive learning such as the basolateral amygdala ^21,22^. Yet, the role of inhibition in cortical circuits shaping specialized neuronal networks during associative learning is still poorly understood. A compelling hypothesis is that local inhibition could modulate feedback inputs received by sensory neurons from higher-order brain areas ^19,20,23^.

To investigate the role of cortical inhibition in associative learning, we recorded neuronal activity using chronic extracellular electrophysiology as mice were trained on a Go/No-go auditory discrimination task. We focused on SST+ neurons because they provide widespread and dense inhibition in the cortex ^24,25^, control the apical tuft dendrites of excitatory neurons ^24,26,27^, and are suggested to gate sensory learning ^28^. We show that, unlike other neurons, SST+ neurons reduce their response to sounds during hit trials. Using optogenetic manipulation, we also demonstrate that a reduced activity of SST+ neurons facilitates associative learning. Since silencing SST+ neurons does not improve bottom-up signal coding, we hypothesized that SST+ neurons could gate top-down inputs involved in associative learning. As top-down inputs, we focused on the axons from the orbitofrontal cortex (OFC) because this higher-order region is involved in outcome-expectancy based decision making ^23^ and has been shown to influence sensory learning ^19^. Using dual-optogenetic circuit mapping combined with extracellular electrophysiological recordings, we show that SST+ neurons control inputs from the OFC. Finally, we provide evidence that the OFC inputs controlled by SST+ neurons are essential for efficient auditory associative learning. Overall, our results indicate that SST+ neurons in the auditory cortex play a key role in the integration of bottom-up and top-down signals during associative learning.

## Results

### Associative learning increases neuronal sound discriminability

We first assessed the plastic changes occurring in primary sensory cortices during associative learning. Previous studies have shown that A1 is necessary for complex, but not simple, auditory discrimination tasks ^29–31^. For example, it is required to distinguish between a pure-frequency tone and a frequency-modulated sweep ^29^. Therefore, we trained head-fixed mice to discriminate between these sounds (Figure 1A). Mice were trained to lick a spout after the Go sound to receive a reward and to withhold licking after the No-go sound to avoid a mild air-puff (Figure S1A, B). Their performance progressively improved (Figure 1B), reaching the high performance criterion (80% of correct responses) on average within six days (Figure S1C). Mice maintained a high hit rate but showed a significant reduction in the false alarm rate when comparing the first day of auditory discrimination training (early stage) to the session where they achieved 80% of correct responses (expert stage; Figure S1D). Response latency to both Go (hit trials) and No-go tones (false alarm trials) remained unchanged throughout the learning process (Figure S1E).

**Figure 1.**
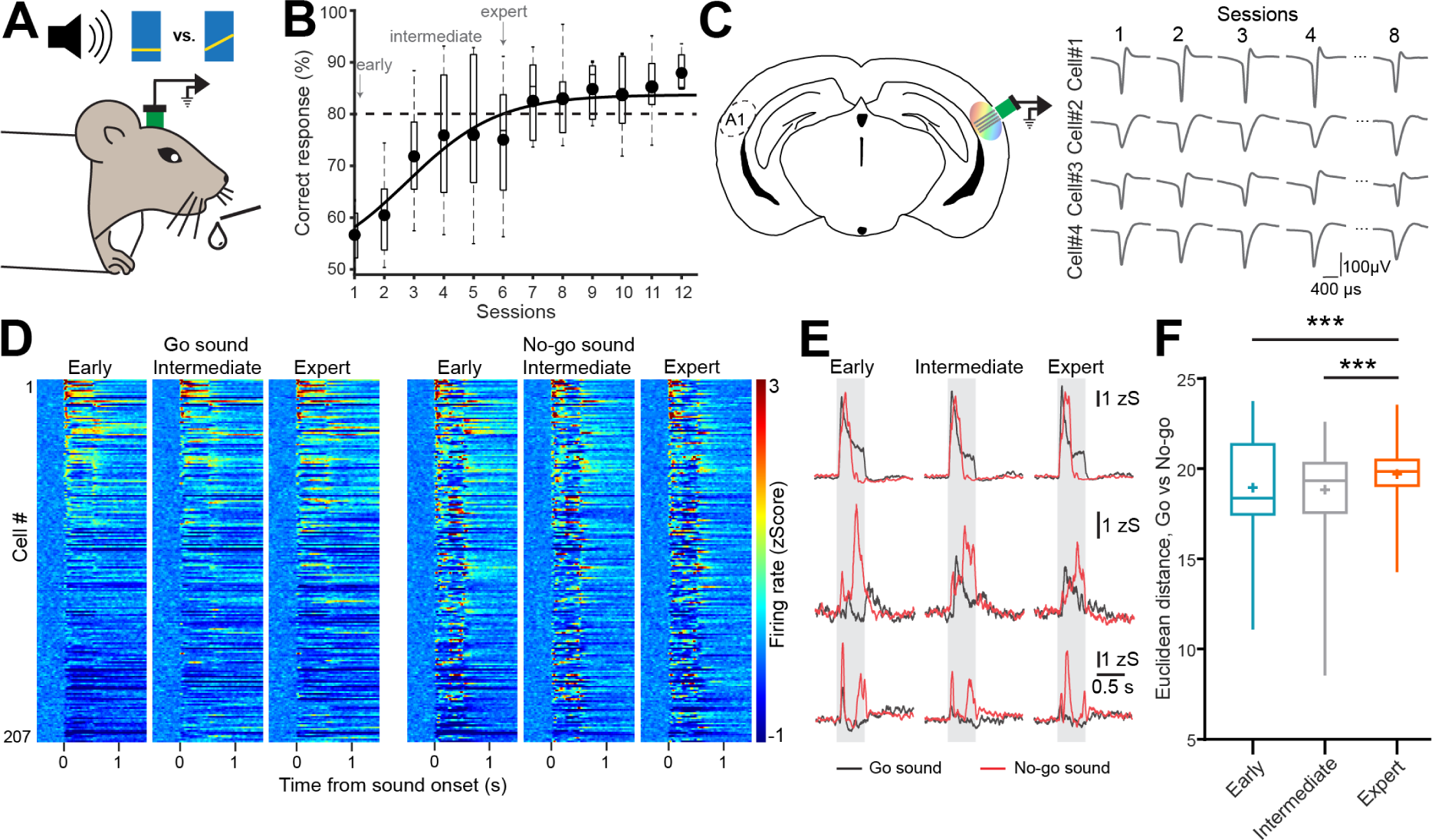
Associative learning increases neuronal sound discriminability. **(A)** Schematic of a head-fixed mouse learning a Go/No-go auditory discrimination task. **(B)** Average performance curve of mice learning an auditory Go/No-go discrimination task (*n*=10). **(C)** Schematic of chronic extracellular recordings in A1 (left). Example waveforms of action potentials from four neurons recorded in A1 over multiple sessions (right). **(D)** Peristimulus time histogram (PSTH) of Go (left) and No-go (right) sound evoked responses of A1 neurons (n=207) at three stages of learning. Heatmaps are sorted based on descending firing rate for Go sound at early stage. **(E)** Mean sound-evoked responses for Go (black) and No-go sounds (red) at three stages of learning for three example neurons (each row). Gray shaded areas represent sound presentation. **(F)** Euclidean distance of Go vs No-go responses as a function of learning stages. (*n*=5 mice, 207 neurons, RM one-way ANOVA *F*(2, 410)=10.41, *P*<0.0001 with Tukey’s multiple comparison test: early vs. expert: *P*<0.0008, intermediate vs. expert: *P*<0.0001, *** *P*<0.001). Boxplots represent the min, 25^th^ percentile, median, 75^th^ percentile, and max; + represent the mean.

To track changes in neuronal activity during learning, we implanted mice with chronic extracellular electrodes in A1 (Figure 1C). This technique allows for the recording of single-unit activity from the extracellular space over extended periods ranging from weeks to months ^32–35^. We collected neuronal data from each training session and for some units we were able to record their activity over multiple days (see Methods) (Figure 1C). Only sorted units present throughout the entire behavioral training (*n*=207, Figure 1D) were included in the analysis. This enabled us to systematically monitor changes in each neuron’s responses to the Go and the No-go sounds over the course of learning (Figure 1D, E).

We initially hypothesized that learning would reduce variability in neuronal responses to the learned sounds. To test this, we compared the coefficient of variation of neuronal responses to Go and No-go sounds across the early, intermediate (approximately 70% of correct response), and expert stages. Since movement has been shown to reduce neuronal activity in the auditory cortex ^36–38^, we restricted the analysis window to the first 250 ms of sound presentation to minimize any major motor activity related to the behavioral response (Figure S1E). Notably, trial-to-trial variability of sound-evoked activity transiently increased at the intermediate stage for both the Go and No-go sounds (Figure S2A), but remained unchanged between the early and expert stages. We next examined whether this trial-to-trial variability was correlated among simultaneously recorded cells by assessing noise correlation, which reflects shared functional inputs ^39^. Similar to variability, noise correlation increased only at the intermediate stage and remained unchanged between the early and expert stages (Figure S2B, C). Although variability did not decrease with learning as we initially hypothesized, these results suggest that significant functional plasticity occurs during the intermediate stage of learning.

We then aimed to determine whether this plasticity changed the capacity of A1 to encode the Go and No-go sounds. To achieve this, we calculated the Euclidean distance between responses to Go and No-go trials for each neuron. The Euclidean distance increased at expert stage (Figure 1F), indicating that learning enhanced discriminability between the two sounds, consistent with the behavioral performance. Together, our results show that improved discriminability performance is associated with a strengthening of the neuronal discriminability of sounds with different behavioral valence.

### Increased discriminability is associated with a subset of neurons gaining selectivity for the learned sounds

An increased discriminability may relate to a strengthening of the selectivity for one sound or for both. It could also reflect changes affecting all neurons or only part of the neuronal population. To further understand the underlying mechanisms, we computed a selectivity index (SI) for each behavioral stage, defined for each neuron as the difference between the mean responses to the Go and No-go sounds divided by the pooled variability ^8,28^. As expected, neuronal selectivity was strongly heterogeneous at the early stage (Figure 2A). Surprisingly, this heterogeneity persisted at all learning stages as reflected in the mean SI, absolute SI, and change of SI, which did not vary significantly with learning (Figure 2B-D). At each stage, the proportion of neurons showing a preference for one sound remained stable (Figure S3A). Thus, the change of sound discriminability observed at the population level (Figure 1G) did not seem to be driven by an overall increased selectivity at the single cell level.

**Figure 2.**
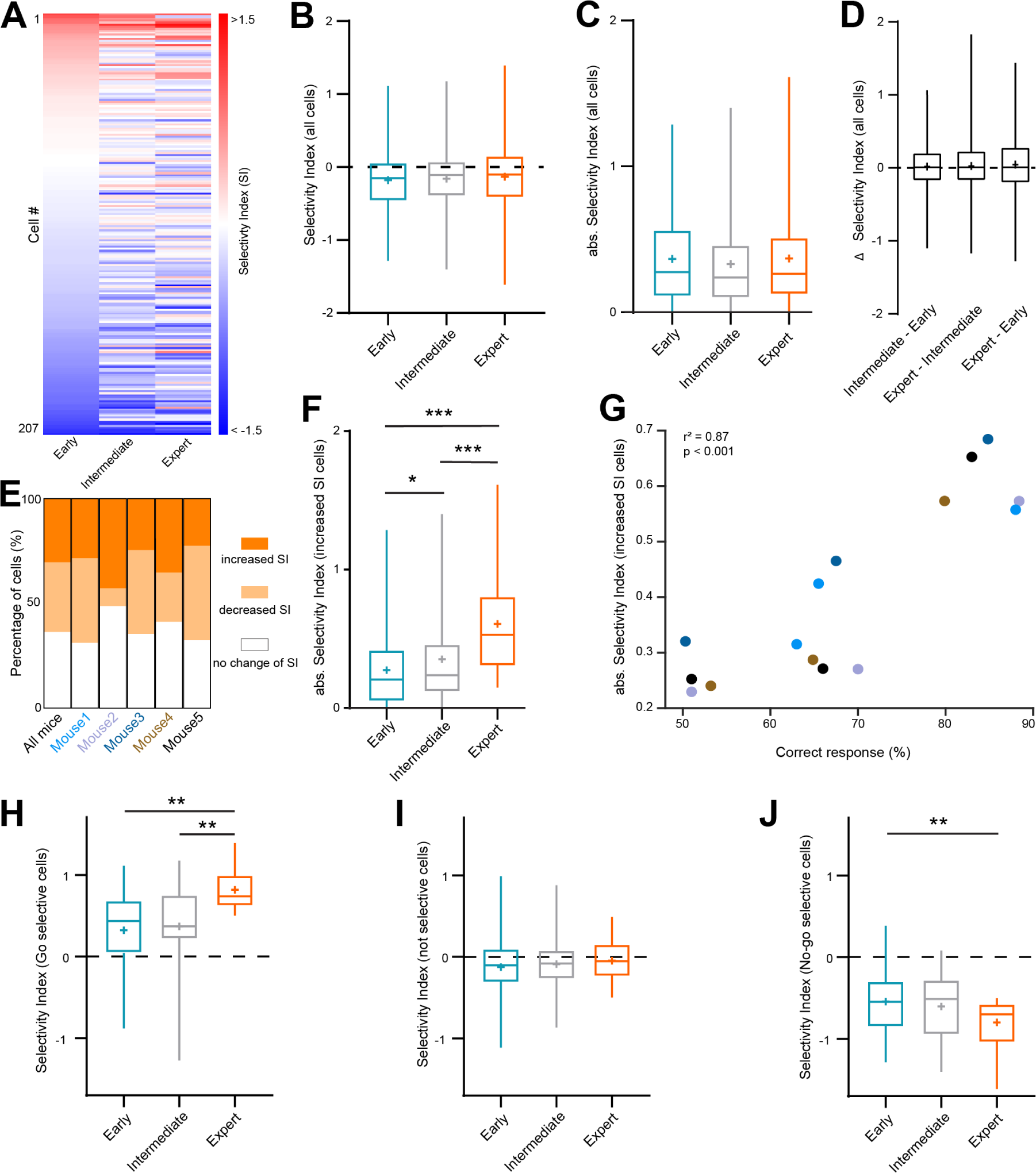
Increased discriminability is associated with a subset of neurons gaining selectivity for the learned sounds. **(A)** Selectivity of neurons for the Go or No-go sound over learning stages. **(B)** Mean SI over learning stages. **(C)** Mean absolute SI over learning stages. **(D)** Change of SI between the different learning stages. **(E)** Percentage of neurons showing significant change of selectivity with learning (from early to expert stage, bootstrap procedure). **(F)** Mean absolute SI of neurons with a significant increased selectivity over learning. (*P*<0.0001, post hocs: early vs. intermediate: *P*=0.0378, early vs. expert: *P*<0.0001, intermediate vs. expert: *P*<0.0001). **(G)** Correlation between the mean absolute SI of neurons with a significant increased selectivity and the behavioral performance for each mouse and each learning stage (Pearson correlation). Each color represents an individual mouse (*n*=5, same as in E). **(H)** SI of neurons selective for the Go sound (SI > 0.5) at expert stage as a function of learning stages. (*P*=0.0007, post hocs: *P*=0.003). **(I)** Same as H for neurons with no preference (-0.5 < SI > 0.5) at expert stage. **(J)** Same as H for neurons with No-go preference (SI < -0.5) at expert stage. (*P*=0.005, post hoc: *P*=0.0041). (*n*=207 neurons, Friedman test with Dunn’s multiple comparison test, * *P*<0.05, ** *P*<0.01, *** *P*<0.001). Boxplots represent the min, 25^th^ percentile, median, 75^th^ percentile, and max; + represent the mean.

This led us to hypothesize that only a fraction of neurons, rather than the entire population, was responsible for the change in discriminability. Among all recorded neurons, 30% (63/207 neurons) increased their selectivity through learning (Figure 2E). Among these neurons, some showed an increased preference for the Go sound, while others favored the No-go sound (Figure S3B, C), suggesting that the enhanced discriminability acquired after learning was not dominated by the increased selectivity for a single sound. Interestingly, these neurons exhibited a progressive strengthening of selectivity throughout learning, with a notable increase after the intermediate stage (Figure 2F). Furthermore, the selectivity of these neurons, averaged for each mouse, strongly correlated with improved behavioral performance (Figure 2G). This correlation did not hold when considering all recorded neurons (Figure S3D), further supporting the specificity of learning-related plasticity within this subset of neurons. We then went back to single-cell level to confirm that the increased selectivity was attributable to neurons selective for either the Go or No-go sound. We clustered neurons based on their selectivity at expert stage into three groups: Go selective (SI > 0.5), No-go selective (SI < -0.5) and not selective (-0.5 < SI > 0.5). The first two groups showed an increasing selectivity with learning (Figure 2H-J). Overall, this data provides evidence that sensory associative learning is driven by a subset of neurons which progressively gain selectivity either for the Go or No-go sound throughout the learning process.

### Somatostatin-expressing interneurons maintain their selectivity for the learned sounds

The progressive increase in selectivity observed during learning could reflect reinforced inputs leading to improved performance. Since local inhibition controls how local networks process sensory inputs ^40^, we next questioned whether changes in selectivity could also be observed in inhibitory neurons. We focused on SST+ interneurons because they target the apical dendrites of excitatory neurons ^41^, modulating their ability to process inputs ^41–43^. More importantly, it has been reported that excitatory neurons in the visual cortex that exhibit the highest anticorrelation with SST+ neurons before associative learning undergo the strongest learning-related changes ^28^. This suggest that SST+ neurons could play a role in sensory associative learning.

We used optotagging ^44^ to identify SST+ neurons as light-activated neurons in mice expressing channelrhodopsin (ChR2) in SST+ neurons (SST-ChR2 mice) (Figure 3A, B). Consistent with our previous analysis, we retained only light-activated neurons that were present throughout the entire learning process (*n*=18) (Figure S4A). In this subpopulation as well, the SI exhibited strong heterogeneity (Figure 3C). The selectivity remained stable throughout learning (Figure 3D, E). Similarly, sound preference at the population level remained stable across learning stages (Figure S4B). 50 % of SST+ neurons (*n*=9, bootstrap procedure) maintained a stable SI during learning (Figure 3G). SST+ neurons characterized by a significant increase in SI (28%, *n*=5, Figure 3G and Figure S4C) developed selectivity for either the Go or No-go sound (Figure S4D). When clustering the SST+ neurons based on their SI at the expert stage, we observed no change during learning (Figure S4E-G), in contrast to SST-neurons (Figure 2H, J). Overall, SST+ neurons did not show a clear change of their selectivity throughout the learning process.

**Figure 3.**
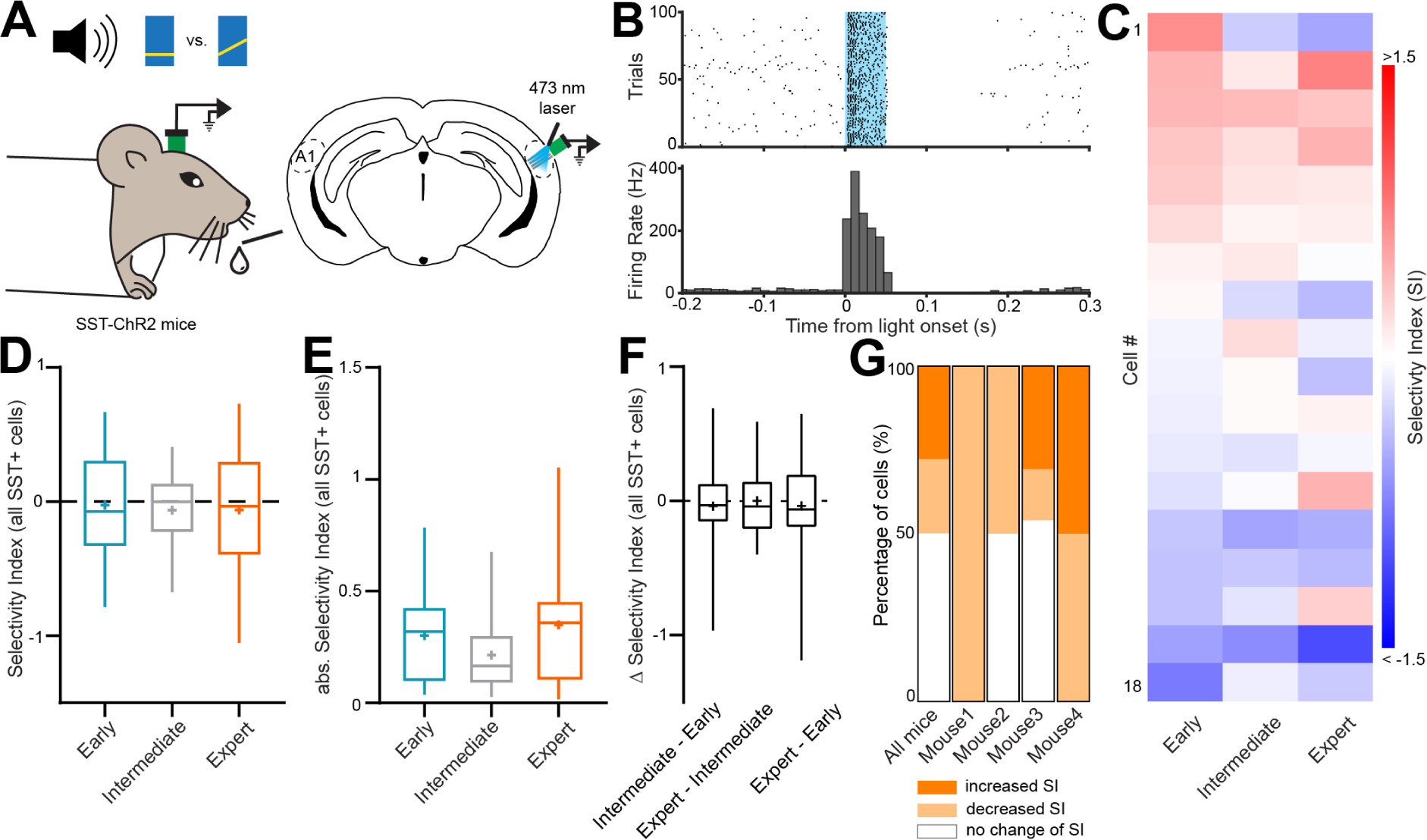
Somatostatin-expressing interneurons maintain their selectivity for the learned sounds. **(A)** Schematic of optogenetic stimulation of SST+ neurons in A1 during chronic extracellular recordings in mice learning a Go/No-go auditory discrimination task. **(B)** Example light-evoked response of a putative SST+ neuron. **(C)** Selectivity of SST+ neurons for the Go or No-go sound over learning stages. **(D)** Mean SI of SST+ neurons over learning stages. **(E)** Mean absolute SI of SST+ neurons over learning stages. **(F)** Change of SI of SST+ neurons between the different learning stages. **(G)** Percentage of SST+ neurons showing significant change of selectivity with learning (bootstrap procedure). (*n*=4 mice, 18 neurons). Boxplots represent the min, 25^th^ percentile, median, 75^th^ percentile, and max; + represent the mean.

### Learning-induced neuronal plasticity is characterized by opposite changes in SST+ and SST-neurons

Since associative learning links an event to its behavioral meaning, we next investigated whether the plasticity of neuronal response we observed with learning were specific to the sounds themselves, or if learning could differentially affect neuronal activity based on the behavioral outcome (i.e. lick or no lick). To this end, we extracted the neuronal activity for each behavioral outcome at each learning stage for optotagged SST+ neurons and non-optotagged SST-neurons (Figure S5). We then computed the difference in sound-evoked responses (difference in peristimulus time histograms, ΔPSTH) between the expert and early stages (Figure 4A) to directly observe the influence of learning on neuronal activity. This approach allowed us to directly compare learning-driven neuronal activity for SST+ and SST-neurons across the different behavioral choice-related trials (Figure 4B-D).

**Figure 4.**
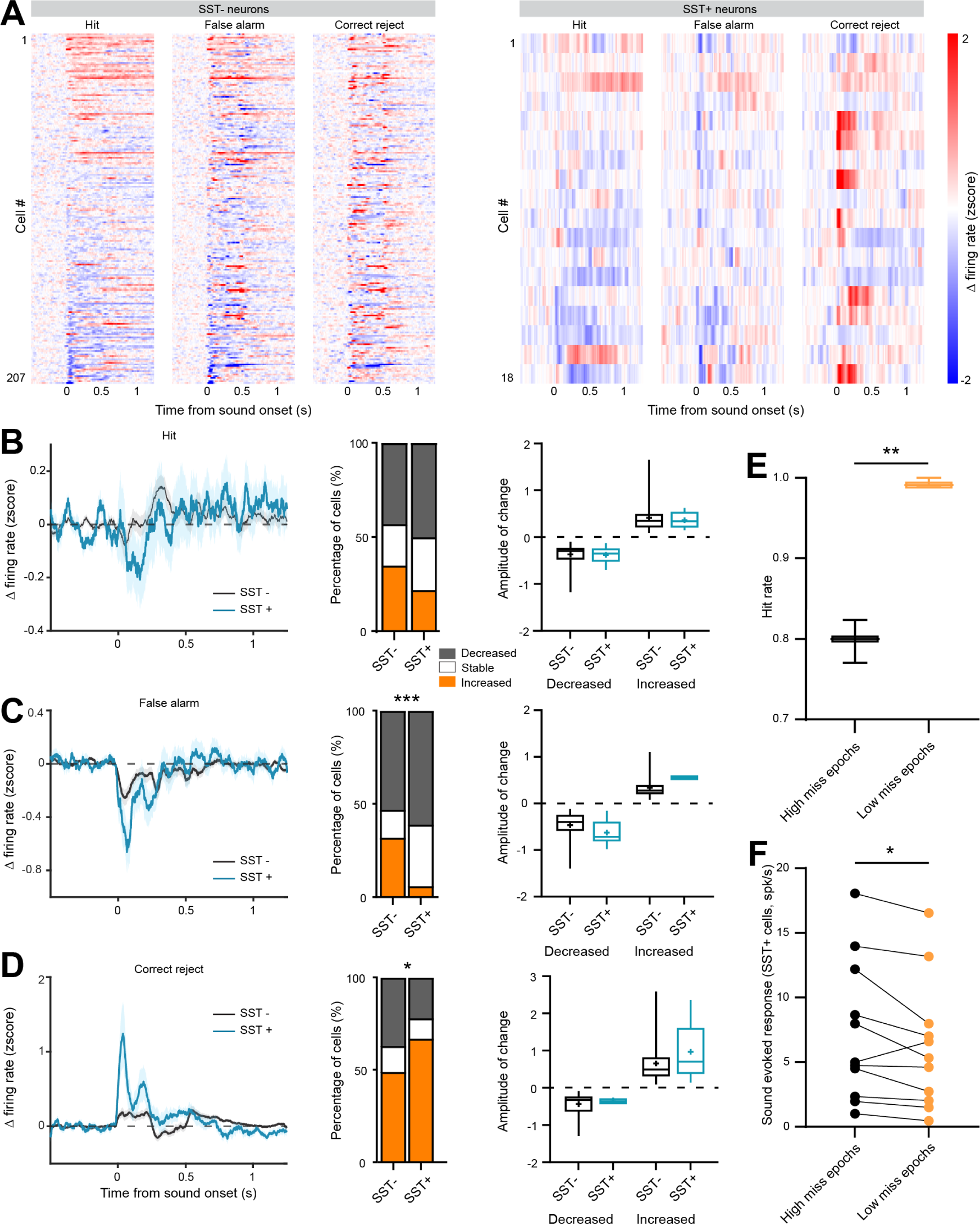
Learning-induced neuronal plasticity is characterized by opposite changes in SST+ and SST-neurons. **(A)** ΔPSTH between the early stage and the expert stage for each neuron segregated between hit trials, false alarm trials and correct reject trials for non-optotagged SST-neurons (left, 207 neurons) and optotagged SST+ neurons (right, 18 neurons). Heatmaps are sorted based on maximum change of response in hit trials. Sounds last 0.5s. **(B)** Average ΔPSTH (left), percentage of neurons with changed response (middle) and amplitude of response’s change (right) for hit trials (SST-*n*=207, SST+ *n*=18). **(C)** Same as B for false alarm trials. (ꭓ^2^ test, *P*<0.0001). **(D)** Same as B for correct reject trials. (ꭓ^2^ test, *P*=0.0307). **(E)** Hit rate of selected epochs. (paired t-test, *P*=0.0044). **(F)** Sound-evoked responses of SST+ neurons as a function of behavioral performance. (*n*=11, Mann-Whitney test, *P*=0.0186). (* *P*<0.05, ** *P*<0.01, *** *P*<0.001). Boxplots represent the min, 25^th^ percentile, median, 75^th^ percentile, and max; + represent the mean.

For the rewarded sound, the low miss rate (Figure S1D) prevented accurate analysis of the neuronal activity specifically associated with miss trials. In hit trials, SST-neurons showed a stable response with learning, whereas SST+ neurons surprisingly exhibited a decreased response (Figure 4B, left). We questioned whether the changed responses were due to an overall modified response of all neurons or rather to a shift in the number of neurons responsive to the Go sound. Half of the SST+ neurons showed a decreased response in hit trials, while a quarter exhibited an increased response. For SST-neurons, only a third showed a decreased response, and another third showed an increased response (Figure 4B, middle). Moreover, the amplitude of change was equivalent for both SST+ and SST-neurons (Figure 4B, right). Thus, the decreased response observed in SST+ neurons was driven by a high number of SST+ cells with a decreased response rather than by the strength of response’s change. Overall, these findings indicate that, unlike for SST-neurons, learning decreases the activity of SST+ neurons for the rewarded sound.

For the non-rewarded sound, we could separate the data into false alarm (FA) and correct reject (CR) trials (Figure 4C, D). Here, we observed that SST+ and SST-neurons displayed similar changes associated with learning: a decreased response during FA trials (Figure 4C) and an increased response during CR trials (Figure 4D). Thus, learning affected the responses of SST+ and SST-neurons to the non-rewarded sound in the same manner. Notably, the reduced response of SST+ neurons associated with learning during FA trials was not paralleled with an increased response of SST-neurons, as was seen for hit trials. This may suggest the involvement of different microcircuits for interpreting rewarded and non-rewarded sensory stimuli.

Interestingly, the changes occurring between the early and intermediate stages were similar to those observed between the early and expert stages (Figure S6): a decreased response of SST+ neurons and an increased response of SST-neurons in hit trials, along with comparable changes for SST+ and SST-neurons during FA and CR trials. This further confirms the specificity of changes observed during learning for each behavioral outcome.

Next, we asked if reduced activity of SST+ neurons in response to the Go sound leads to improved performance. We grouped epochs according to hit rate (Figure 4E) and analyzed the associated sound-evoked activity in SST+ neurons. SST+ neurons showed a decreased response in epochs characterized by higher hit rates (Figure 4F). This supports the idea that a reduced activity in SST+ neurons facilitates better discrimination performance.

### Reduced activity of SST+ neurons facilitates associative learning

We next tested whether the reduced activity of SST+ neurons during Hit trials supports associative learning. We used optogenetics to activate SST+ neurons and counteract their learning-associated reduction in activity (Figure 4B). We applied blue light bilaterally over A1 in mice expressing ChR2 in SST+ neurons (Figure S7C) during the Go/No-go discrimination task (see Methods) (Figure 5A). The light was turned on during the response window, i.e. two seconds from sound onset. The results show that the learning curve of SST-ChR2 mice was shifted to the right when compared to control mice that did not express opsins (Figure 5B), suggesting a reduced capacity to learn the association between sounds and their meaning. Indeed, mice needed significantly more time to learn the task (Figure 5C). However, no diffierence in performance or latencies for both hit and false alarm responses were observed between groups at the end of training (Figure 5D, E), suggesting that although SST+ neuron activation drastically impaired the learning process, performance at the expert stage was similar. These results underscore the importance of a reduced activity of SST+ neurons for efficient associative learning. To further investigate the effiect of SST+ neurons photoactivation on the neuronal network, we performed acute extracellular recordings in awake head-fixed naive mice listening to the Go and No-go sounds. The results confirmed that activation of inhibitory SST+ neurons drastically decreased the activity of surrounding neurons (Figure S8A, B). This activation also reduced sound discriminability, as indicated by decreased Euclidean and Cosine distances between trial-to-trial response vectors for the Go and the No-go sounds. This was further supported by an increased correlation between Go and No-go responses (Figure S8C-E). The diminished discriminability aligns with the slower learning speed at the behavioral level when photoactivating SST+ neurons.

**Figure 5.**
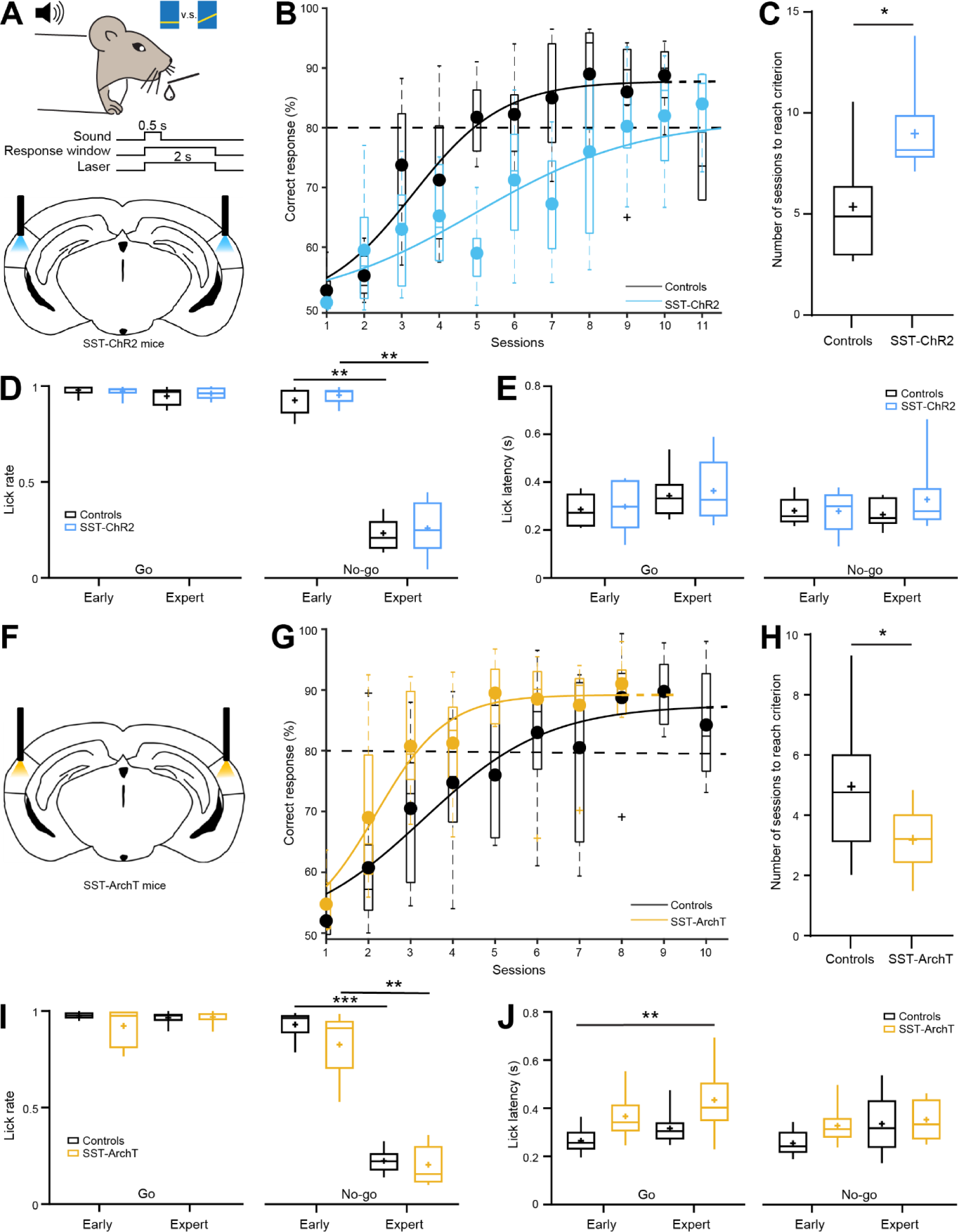
Reduced activity of SST+ neurons facilitates associative learning. **(A)** Schematic of bilateral optogenetic stimulation of SST+ neurons in the auditory cortex of SST-ChR2 mice learning an auditory Go/No-go discrimination task. **(B)** Average learning curve of the Go/No-go task with blue light illumination in SST-ChR2 mice (*n*=6) and SST-Cre control mice (*n*=7). **(C)** Average number of sessions to reached 80% of correct responses (Mann-Whitney test, *P*=0.0221). **(D)** Comparison of the lick rate in response to the Go sound (left) and No-go sound (right) at early and expert stage for SST-ChR2 and control mice (Kruskal-Wallis test, *P*=0.0003, Dunn’s multiple comparison test, *P*=0.0078 and 0.0056). **(E)** Comparison of the lick latency in response to the Go sound (left) and the No-go sound (right) at early and expert stage for SST-ChR2 and control mice. **(F-J)** Same as A-E for the comparison between SST-ArchT mice (*n*=9) and control mice (*n*=10). (H, unpaired t-test, *P*=0.047. I, Kruskal-Wallis test, *P*<0.001, Dunn’s multiple comparison test, *P*=0.0003 and 0.005. J, Kruskal-Wallis test, *P*=0.0043, Dunn’s multiple comparison test, *P*=0.0034). Boxplots represent the min, 25^th^ percentile, median, 75^th^ percentile, and max; + represent the mean.

If the reduced activity of SST+ neurons observed has a behavioral relevance, we hypothesized that further reducing it should facilitate associative learning. To test this, we applied yellow light bilaterally over A1 in mice expressing the inhibitory opsin archaerhodopsin (ArchT) in SST+ neurons (Figure S7d) during the Go/No-go discrimination task (Figure 5F). Optogenetic silencing SST+ neurons shifted the learning curve of SST-ArchT mice to the left (Figure 5G), indicating that these mice reached high performance faster than control mice (Figure 5H). As before, no diffierences in performance or latencies for hit and false alarm responses were observed between groups at the end of training (Figure 5I, J), suggesting that optogenetic modulation did not introduce non-specific factors affiecting behavior.

These results demonstrate that reduced SST+ neuron activity, as observed in our electrophysiological recordings (Figure 4B), facilitates sensory associative learning. Reduced activation of SST+ neurons may drive increased sensory responses triggered by bottom-up inputs ^45^. This mechanism could enhance the saliency of sensory signals, thereby facilitating sensory learning. To test this hypothesis and better understand the control exercised by SST+ neurons on the neuronal network during bottom-up signal processing, we recorded Go and No-go sound-evoked responses in A1 of awake head-fixed naïve mice. Photoinhibition of SST+ neurons increased the sustained activity of neighboring neurons (Figure S8F, G) but did not enhance the sound onset response (0-50 ms from sound onset). Surprisingly, it even decreased sound discriminability (Figure S8H-J) by non-specifically increasing overall neuronal activity in A1 during both sound presentations (Figure S8F).

Overall, our data show that decreased SST+ neuron activity facilitates associative learning. Since silencing SST+ neurons does not improve sound discriminability, this suggests that the facilitation does not result from enhanced bottom-up signaling but could rather come from top-down input modulation.

### SST+ neurons control higher-order inputs from the orbitofrontal cortex to A1 neurons

Top-down input from higher-order brain areas to primary sensory cortices is crucial for associative learning ^20,23^. Increasing evidence suggests that the OFC is involved in outcome-expectancy decision making ^23,46^ and sends information to A1 ^47,48^. The direct input from OFC to A1 neurons was confirmed through anterograde and retrograde tracing, as well as *in vitro* patch-clamp recording of blue light evoked post-synaptic currents in A1 neurons when OFC neurons expressed ChR2 (Figure S9). We hypothesized that the reduced activity of SST+ neurons could facilitate local integration of top-down inputs from OFC. To test this, we expressed ChR2 in OFC neurons of SST-ArchT mice and recorded neuronal activity in A1 using *in vivo* extracellular electrophysiology (Figure 6A). By applying blue light over A1, we stimulated axons of OFC neurons and observed that 72% (174/240 neurons) of auditory neurons responded OFC axon stimulation (Figure S10A). These neurons were considered as receiving input from the OFC (see Methods) and were included in further analysis. In 50% of trials, we applied yellow light to downregulate SST+ neuron activity, simultaneously with blue light (Figure 6B). At the population level, reducing SST+ neuron activity did not alter the response of auditory neurons to OFC axon stimulation (Figure S10B). We then characterized the change of light-evoked response for each single neuron (Figure 6C). Silencing SST+ neurons increased the response to OFC input stimulation in 13% of neurons, decreased it in 26%, and had no effiect in 61% (Figure 6D-F). The strength of SST+-mediated modulation of OFC inputs was similar for neurons with an increased response than for neurons with a decreased one (Figure 6G). This indicates that SST+ neurons have the capacity to control the input from OFC received by a substantial portion of auditory neurons.

**Figure 6.**
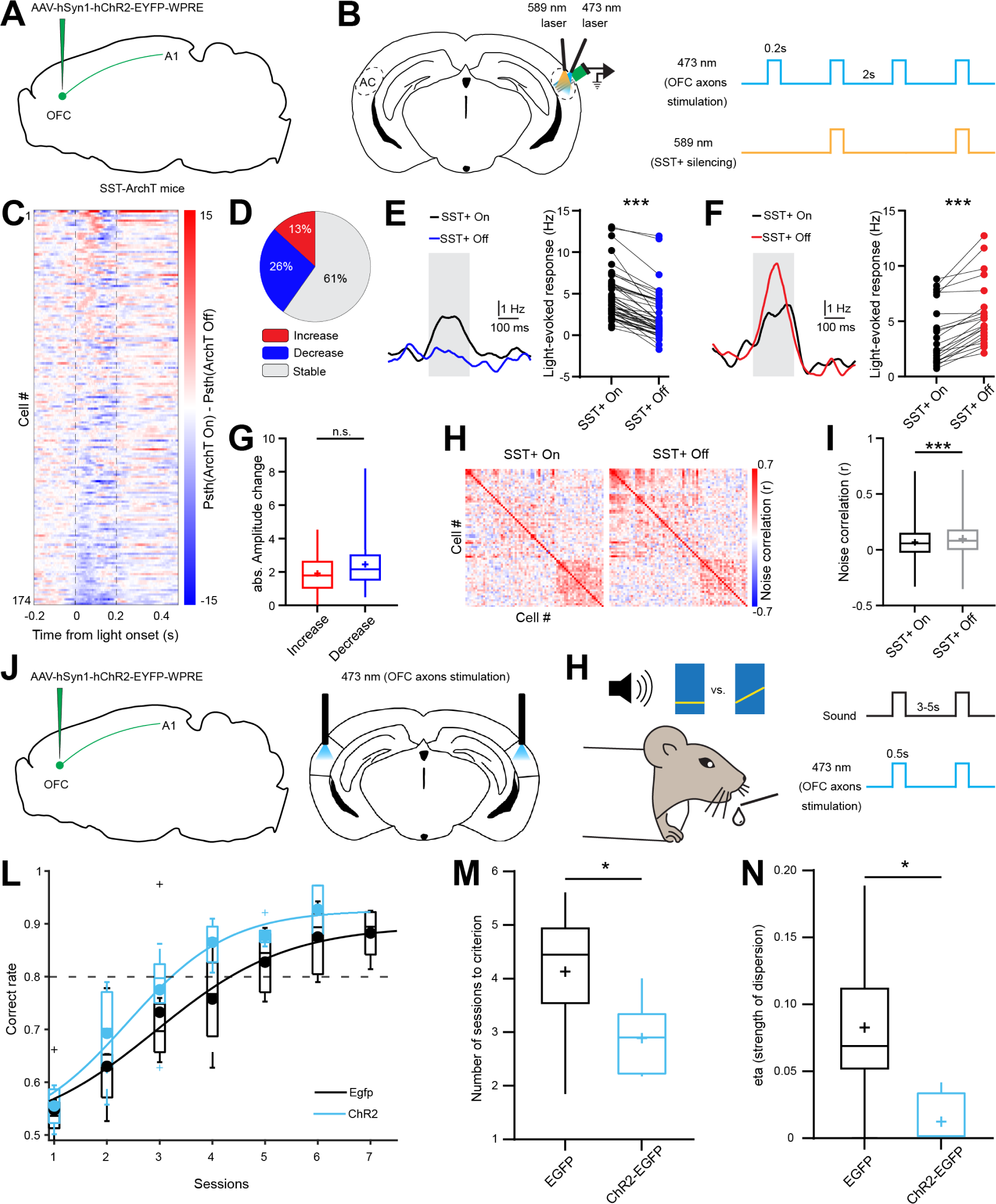
SST+ neurons control inputs from OFC to A1 involved in associative learning. **(A)** Schematic of the procedure to express ChR2 in OFC neurons in SST-ArchT mice. **(B)** Schematic of light-evoked responses recordings using extracellular recordings in A1 triggered by optogenetic stimulation of axons of OFC (left). Protocol of dual optogenetic stimulation of OFC axons expressing ChR2 with blue light (473 nm, 100% trials) and optogenetic silencing of SST+ neurons (589 nm, 50% trials) (right). **(C)** Diffierence of PSTH of blue light-responsive neurons with and without SST+ neurons silencing. **(D)** Percentage of neurons with a significant change of response to blue-light stimulation when SST+ neurons are silenced (bootstrap procedure). **(E)** Example neuron with a decreased response to blue-light stimulation when SST+ neurons are silenced (left). Mean blue-light response for neurons with a decreased response with SST+ neurons silencing (n= 45, Wilcoxon test*, P*<0.0001). **(F)** Example neuron with an increased response to blue-light stimulation when SST+ neurons are silenced (left). Mean blue-light response for neurons with an increased response with SST+ neurons silencing (*n*=22, Wilcoxon test, *P*<0.0001). **(G)** Absolute change of the amplitude of light-evoked response of neurons with a significant change of response to blue-light stimulation when SST+ neurons are silenced (n=45 and 22, Mann-Whitney test, *P*=0.13). **(H)** Example noise correlation matrices from the same mice with or without optogenetic silencing of SST+ neurons. **(I)** Average noise correlation with or without optogenetic silencing of SST+ neurons (*n*=4266, paired t-test, *P*<0.0001). **(J)** Schematic of the procedure to express ChR2 in the axons of OFC neurons and their bilateral optogenetic stimulation over A1 using blue light (473 nm). **(K)** Protocol of stimulation of OFC axons over the learning of a Go/No-go auditory discrimination task. **(L)** Average learning curve of the Go/No-go task with blue light illumination in Egfp-ChR2 mice (*n*=6) and Egfp control mice (*n*=7). **(M)** Average number of sessions to reach 80% of correct responses (unpaired t-test, *P*=0.0492). **(N)** Average eta factor of fit function (unpaired t-test, *P*=0.0189). Boxplots represent the min, 25^th^ percentile, median, 75^th^ percentile, and max; + represent the mean.

We next aimed to determine whether the control exerted by SST+ neurons over the ability of A1 neurons to process inputs from OFC is functionally relevant. Since noise correlation is thought to reflect shared input among neurons ^39^, we compared the ability of OFC inputs to trigger correlated activity in A1 in the presence or absence of SST+ modulation (Figure 6H). We observed a significant increase in noise correlation when SST+ neurons were silenced (Figure 6I), suggesting that reduced SST+ neuron activity facilitates the processing of inputs from OFC. We then tested whether inputs from OFC to A1 facilitate associative learning. We expressed ChR2 bilaterally in OFC and applied blue light bilaterally over A1 to stimulate OFC axons (Figure 6J) during the Go/No-go discrimination task (Figure 6K). The learning curve of OFC-ChR2 mice was shifted to the left compared to controls (OFC-EGFP mice) (Figure 6L), indicating that stimulating OFC inputs to A1 accelerated learning (Figure 6M). Additionally, it reduced variability in learning among mice, as shown by a reduced eta factor, which reflects the dispersion of data points to the fit function (Figure 6N). This suggests that stimulating OFC inputs led to a more efficient performance improvement across training sessions. Lick rates and latencies were similar between OFC-ChR2 and OFC-EGFP groups (Figure S10C, D), indicating no effiect of OFC axons stimulation on perception and motivation. Finally, to assess whether OFC axons stimulation could alter the perception of sensory cues, we tested trained mice on a Go/No-go perceptual task in which the Go cue was kept constant while the No-go cue frequency content was altered, making the No-go cue more similar to the Go cue, increasing the task difficulty (Figure S10E). Activation of OFC axons over A1 did not change the mice’s perceptual capacity (Figure S10E).

In summary, our data demonstrates that SST+ neurons control inputs from OFC which are necessary for associative learning. Moreover, we provide evidence that the influence of OFC on A1 during associative learning does not alter sensory processing, suggesting that it is rather related to the outcome expectancy or valence of the stimulus ^23^.

## Discussion

Understanding how bottom-up and top-down information converge to drive sensory learning is a central question in neuroscience. Here, we show that increased signal discriminability is driven by a subset of neurons that gain selectivity for learned stimuli. During learning, SST+ neurons decrease their response during hit trials. This decrease improves learning by facilitating inputs from the OFC. Overall, our data demonstrates for the first time that cortical SST+ neurons gate the interaction between sensory bottom-up and outcome-expectancy top-down information necessary for associative learning.

We report that decreased activity of SST+ neurons in A1 facilitates associative learning. Several mechanisms could explain this association. First, SST+ neurons are known to drive network suppression ^25^. They could therefore provide inhibition over a wide area thanks to the widespread of their axonal arborization ^25,49^ and contact a large number of surrounding excitatory neurons ^24^. This structural characteristic allows them to efficiently suppress neuronal activity in response to non-specific sensory signals, thereby enhancing sensory coding ^25^. Therefore, a decreased activity of SST+ neurons would theoretically alter sensory coding. Indeed, we show that photoinhibition of SST+ neurons reduces sound discriminability at the neuronal level. This seems contradictory to the increased selectivity of SST-neurons that we observe during learning. Moreover, we show that the decreased activity of SST+ neurons is correlated with enhanced performance. As a second plausible mechanism, we propose that SST+ neurons predominantly inhibit excitatory neurons on their apical dendrites ^26,27^, where these neurons receive input from higher-order brain areas ^12^. Consequently, reduced inhibition from SST+ neurons could locally increase the excitability of excitatory neurons, promoting the reinforcement of top-down inputs coding for the valence of the behavioral outcome. Our data support this hypothesis: we show that SST+ neurons gate direct inputs from the OFC that are involved in associative learning.

Higher-order brain areas also show increased responses to learned sensory cues ^5,50^. The feedback they provide to sensory areas is suggested to be essential for associative learning ^9,20,46,47^. The OFC is implicated in outcome-based decision making ^23^, and several studies have reported that the OFC sends contextual and outcome-related information to sensory areas ^19,23,47^, which is critical for building and updating internal models of cue relevance^23,51^. We show that direct input from OFC is present in A1 and is essential for appetitive associative learning. Moreover, we argue that SST+ neurons control this input and that a decreased SST+ neuron activity enhances feedback from the OFC to A1. However, a recent study contradicts these results, showing that activating SST+ neurons in the visual cortex facilitates associative learning of a visual Go/No-go task ^19^. In that study, the authors observed that the OFC sends strong inputs directly to SST+ neurons, leading to V1 inhibition specifically during No-go trials, thereby suppressing cortical activity in response to irrelevant sensory cues. They also found that inhibiting OFC inputs to V1 during No-go trials facilitated learning, but had no effiect during Go trials. Importantly, they did not manipulate SST+ neurons during Go trials. The discrepancies between their findings and ours regarding the effiects of SST+ activation and OFC inputs inhibition on sensory learning could be due to several factors. First, the circuits involved may diffier between sensory areas. Second, in our study we manipulated OFC inputs or SST+ neurons during both Go and No-go sounds to avoid potential behavioral bias caused by light stimulation itself. This strategy might balance diffierential effiects on rewarded (Go) versus non-rewarded (No-go) trials. Indeed, we observed that SST+ neurons enhance OFC inputs for 13% of auditory neurons but reduce them for 25%. This suggests that within the auditory network, SST+ neurons can both enhance and reduce the influence of OFC on auditory neurons, potentially favoring OFC inputs during Go trials and suppressing them during No-go trials.

The analysis of choice-related activity revealed a learning-associated decrease in SST+ neuron activity during hit trials, but an increased activity for correct reject trials, both associated with opposite effiects on the activity of SST-neurons. This suggests that learning drives distinct plasticity depending on trial outcome. When the sound is associated with a reward, SST+ neurons reduce their activity to favor inputs from higher-order brain areas such as the OFC, carrying information about the valence of the outcome. This might be essential for taking the decision to lick and collect the reward. Indeed, during false alarm trials, SST+ neurons also decreased their activity, and the mice do not refrain from licking. In contrast, increased SST+ neuron activity during correct reject trials, may help suppress bottom-up signals, preventing the mice from licking in response to the No-go sound. Overall, this suggests that SST+ neurons are progressively downregulated by the Go sound and upregulated by the No-go sound. How SST+ neurons are diffierentially embedded in neuronal circuits involved in the modulation of feedback related to cues with positive or negative valence remains to be addressed.

Taken together, our results highlight the critical role of SST+ neurons in gating inputs from higher-order brain areas. Additionally, we provide evidence that OFC inputs to primary sensory areas facilitate associative learning. These inputs are crucial for associating sounds with their behavioral meaning and their reinforcement allows sensory learning.

## Acknowledgements

We thank Pauline Brunet and Philip Williams for technical help and the members of the Brain and Sound Lab for fruitful discussions. We thank Morgane Roth and Andreas Keller for their comments on the manuscript. This work was supported by a Marie Sklodowska-Curie Individual Fellowship (MSCA-IF EU project 894719 – AudiLearn, F.S.), and by the Swiss National Science Foundation (ERC Transfer grant CRETP3-166735 and project grant 310030_197850, T.R.B).

## Author contributions

F.S. performed most of the experiments and analyzed the data. T.Z. performed intracellular recordings. F.S. and T.R.B. designed the study, interpreted the data ad wrote the manuscript with input from T.Z.

### STAR Methods

#### Experimental model and subject details

All experimental procedures were performed in accordance with the University of Basel animal care and use guidelines and were approved by the Veterinary Office of the Canton Basel-Stadt, Switzerland. We used C57BL/6J mice (Janvier Laboratories, France), SST-Cre knock-in line with C57BL/6J background (JAX stock number 013044, Jackson Laboratories, ME, USA), mice derived from crossing a SST-Cre knock-in line with a ChR2-floxed Ai32 line (JAX stock n. 024109, C57BL/6J background) or ArchT-floxed Ai35D line (JAX stock n. 012735, C57BL/6J background). Mice were a mixture of males and females aging from 5 to 12 weeks old from the time of surgery to the end of experiments. Water and food were given *ad libitum*, except for behavioral experiments where food intake was manually regulated by the experimenter starting from the day before the start of the training phase. The animals’ weight was checked daily and maintained above 80% of the weight before food restriction. Mice were single-housed under a 12:12 h light/dark cycle. Experiments were performed in the light phase.

## Method Details

### Surgical procedures

#### Chronic electrode implantation

Mice were anesthetized with a mixture of ketamine/xylazine (80 and 16 mg/kg, respectively) and local analgesia was provided with a subcutaneous injection of bupivacaine/lidocaine (0.01 mg/animal and 0.04 mg/animal, respectively). Ketamine (45 mg/kg) was supplemented during surgery as needed. Perioperative analgesia was provided by injection of buprenorphine (0.1mg/kg, i.p.). Body temperature was maintained at 37°C via a heating pad (FHC, ME, USA) and lubricant ophthalmic ointment was applied on both eyes. A custom-made metal head plate and a ground screw were attached to the skull with super glue (Loctite, Henkel, Germany). For A1 recordings, a craniotomy was performed above the right auditory cortex (ACx) and the brain was covered by silicon oil to prevent drying. A chronic multi-channel extracellular electrode (A4×8-3mm-50-200-177-CM32 or A4×16-3mm-50-200-177-Z64, Neuronexus, MI, USA) was inserted with a motorized stereotaxic micromanipulator (DMA-1511, Narishige, Japan) to determine the location of A1 based on functional tonotopy (caudo-rostral increase in best frequency). An optic fiber cannula (CFMLC21L05, Thorlabs, NJ, USA) was placed next to the electrode over A1. Once the desired location was reached, a silicone casting compound (Kwik-Sil, WPI, FL, USA) was applied over the craniotomy to cover the brain surface before affixing the electrode to the skull with dental cement (Super Bond, C&B, Japan).

#### Acute electrophysiology

Mice were anesthetized with isoflurane (4% for induction and 1.5 to 2.5% for maintenance) and local analgesia was provided with subcutaneous injection of bupivacaine/lidocaine (0.01 mg/animal and 0.04 mg/animal, respectively). Body temperature was maintained at 37°C via a heating pad (FHC, ME, USA) and lubricant ophthalmic ointment was applied on both eyes. A custom-made metal head plate was attached to the skull with super glue (Henkel, Loctite, Germany). A craniotomy was performed above ACx and the brain was covered by silicone oil and silicone casting compound (Kwik-Cast, WPI, FL, USA) during the 2 hour recovery period from the anesthesia to prevent it from drying.

#### Optogenetics

Mice were anesthetized with isoflurane (4% induction and 1.5-2.5% for maintenance), placed in a stereotaxic frame (David Kopf Instruments, CA, USA) and local analgesia was provided with subcutaneous injection of bupivacaine/lidocaine (0.01 mg/animal and 0.04 mg/animal, respectively). Perioperative analgesia was provided by injection of buprenorphine (0.1 mg/kg, i.p.). Body temperature was maintained at 37°C via a heating pad (FHC, ME, USA) and lubricant ophthalmic ointment was applied on both eyes. A custom-made metal head plate was attached to the skull with super glue (Loctite, Henkel, Germany). To manipulate SST neurons in A1, craniotomies were performed bilaterally over ACx in SST-ChR2 mice, SST-ArchT mice, or SST-Cre mice. Optic fiber cannulae (CFMLC21L02, Thorlabs, NJ, USA) were approached to the brain surface, silicone casting compound (Kwik-Sil, WPI, FL, USA) was applied over craniotomies before soldering optic fiber cannulae to the skull with dental cement (Super Bond, C&B, Japan). To visualize or manipulate OFC axons projecting to A1, we injected AAV-1/2-hSyn1-hChR2(H134R)-EYFP-WPRE or AAV-1/2-hSyn1-EYFP-WPRE (200 nL, 40 nL/min) in OFC (AP, 2.5 mm; ML, 1.1 mm; DV, 1.9 mm) with a glass micropipette controlled by a nanoinjector (Nanoliter 2020, WPI, FL, USA). Then craniotomies were performed bilaterally over ACx, optic fiber cannulae (CFMLC21L02, Thorlabs, NJ, USA) were approached to the brain surface, silicone casting compound (Kwik-Sil, WPI, FL, USA) was applied over craniotomies before affixing optic fiber cannulae to the skull with dental cement (Super Bond, C&B, Japan). To visualize long range input to A1, we injected AAV-retro/2-hSyn1-chl-mCherry_2A_iCre-WPRE in A1 (AP, -3.9 mm; ML, 5.5 mm; DV, 0.4 mm) (300 nL, 40 nL/min) with a glass micropipette controlled by a nanoinjector (Nanoliter 2020, WPI, FL, USA).

### Behavioral training

Sounds were generated with a digital signal processor (RZ6, Tucker Davis Technologies, FL, USA) at 200 kHz sampling rate and played through a calibrated MF1 speaker (Tucker Davis Technologies, FL, USA) positioned 10 cm from the mouse’s left ear. Stimuli were calibrated with a wide-band ultrasonic acoustic sensor (Model 378C01, PCB Piezotronics, NY, USA). Mice were trained to discriminate between a Go tone (4 kHz Pure Tone) and a No-go tone (Frequency-modulated Sweep, 4-16 kHz) played at 70dB SPL. The sound lasted 500 ms, the response window lasted 2 s from the sound onset and the inter-trial interval varied between 3 and 5 s to avoid prediction of stimulus presentation. Mice were head fixed and placed in a cardboard tube, facing a plastic spout attached to a piezo sensor to detect licking behavior. Before the training started, mice were food deprived for 24h. On the first day of training, mice were placed in a cardboard tube and habituated to head restriction. From the second day on, all trials consisted of Go sound presentation. Mice were trained to lick during the 2 sec long response window to receive a reward consisting of a drop of soy milk (‘hit trial’). No lick was considered as a ‘Miss trial’. When mice reached 80% of hits, the No-go sound was introduced and represented 50% of trials. A lick in response to the No-go sound (‘false alarm’) resulted in a mild air-puffi and an extra time-out of 3 s. No lick in response to the No-go was registered as a correct reject. When comparing the speed of learning among groups, each discrimination session was restricted to 400 trials. For OFC optogenetic manipulation experiments, when mice reached 80% of correct responses, the perceptual threshold was assessed by progressively decreasing the diffierence between the Go and No-go sound using FMS with decreased frequency ranges (4-16kHz, 4-12kHz, 4-8kHz, 4-6kHz, 4-5kHz and 4-4.5kHz).

### Extracellular Electrophysiological recordings

Recordings were performed in head-fixed mice sitting in a cardboard tube in a sound-attenuating chamber. For acute recordings, the silicone casting compound was removed, and the exposed brain was covered with silicon oil. Multi-channel extracellular electrodes (A4×16-5mm-50-200-177-A64, Neuronexus, MI, USA) were slowly lowered into the brain orthogonal to the surface with a motorized stereotaxic micromanipulator (DMA-1511, Narishige, Japan). For both acute and chronic recordings, the analog signal was amplified, digitized and acquired at 24’414 Hz (RZ2 Bioamp processor, Tucker Davis Technologies, FL, USA). For chronic recordings, blocks of consecutive sessions recordings were concatenated. Putative single and multi-units were identified offline using KiloSort 2.0 (CortexLab, UCL, England) followed by manual inspection of spike shape and signal-to-noise ratio, and auto- and cross-correlograms using Phy 2 (CortexLab, UCL, England). Both single and multi-units were retained for analysis. For chronic recordings, we kept for analysis only neurons present throughout the learning process (from early to expert stages) and with a stable waveform across days (across-sessions waveform correlation higher than 0.8).

### Optogenetics

Optical activation of ChR2 was induced by a blue light laser (473 ± 1nm) and optical silencing through ArchT activation was induced by a yellow light laser (589 ± 1nm) (Shanghai Laser & Optics Century Co., China).

For behavioral experiments, we presented light bilaterally through optic fiber cannulas implanted over A1. The light was presented duringthe response window (during 2 s from sound onset) and for all trials from the beginning of the discrimination task training. To activate SST+ neurons in SST-ChR2 mice or OFC axons expressing ChR2 during behavior, we applied pulse trains of blue light (10 mW, 10 Hz). To inhibit SST+ neurons in SST-ArchT mice, we used a single pulse of continuous yellow light (10 mW).

For concomitant optogenetic activation of OFC axons and SST+ neurons silencing during acute electrophysiological recordings, we injected AAV-1/2-hSyn1-hChR2(H134R)-EYFP-WPRE in the right OFC of SST-ArchT mice. 4-6 weeks after the virus injection, we performed acute extracellular recordings in A1 and placed 2 optic fibers over the craniotomy: one connected to a 473 nm laser and the other connected to a 589 nm laser. Blue light illumination was on at every trial (30 mW, 200 ms). Yellow light was activated every second trial (10 mW, 200 ms).

### Histology

Mice were transcardially perfused with PBS (1x) then with 4% PFA. Brains were collected, postfixed in 4% PFA for at least 4h, cryoprotected in 30% glucose and frozen embedded in OCT (Tissue Teck, Sakura Finetek, Japan) on dry ice. Free floating brain slices (50 µm thick) were collected at the cryostat. Slices were exposed to a blocking solution (BSA 2% in PBS-T 0.05%) for 1h. Slices were then exposed to primary antibody overnight at 4°C. The following primary antibodies were used: rabbit anti-SST (1/500, BMA biomedicals, Switzerland), chicken anti-GFP (1/500, Abcam, UK). After extensive washing in PBS, slices were exposed to a secondary antibody for 2h in the dark at room temperature. The following secondary antibodies were used: anti-rabbit 555 (1/500, Life Technologies, CA, USA), anti-chicken 488 (1/500, Jackson Immunoresearch Laboratories, PA, USA). DAPI (1/1000) was used to stain cellular nucleus. Then slices were mounted between a slide and a glass cover slip. Image were acquired at 10x using a slide scanner (Axioscan Z.1, Zeiss, Germany).

### Intracellular electrophysiology

Four to five weeks after virus injection, mice were anesthetized using isoflurane (5%) and decapitated. The brain was rapidly extracted and immersed in an ice-cold slicing solution containing (in mM): 93 N-Methyl-D-glucamin (NMDG), 2.5 KCl, 1.2 NaH2PO4, 30 NaHCO3, 20 HEPES, 25 glucose, 5 sodium ascorbate, 2 thiourea, 3 sodium pyruvate, 10 MgSO4 and 0.5 CaCl2 (titrated to pH 7.3-7.4 with HCl, continuously bubbled with 95% O2 and 5% CO2, osmolarity ≈310 mOsm). Two coronal cuts were performed to isolate the region containing the auditory cortices and the tissue block was quickly placed on a vibratome (Leica VT 1200S) where coronal slices (300 μm thick) were prepared. Based on visual inspection of anatomical landmarks, slices containing the auditory cortex were first incubated in the slicing solution for 15 minutes at 31 to 33 °C, and subsequently transferred for 45 minutes at room temperature in standard aCSF containing (in mM): 125 NaCl, 2.5 KCl, 25 NaHCO3, 1.25 NaH2PO4, 10 Glucose, 2 CaCl2 and 0.5 MgCl2 (continuously bubbled with 95% O2 and 5% CO2; pH = 7.3, osmolarity ≈ 310 mOsm).

Whole-cell patch clamp recordings were conducted at 31 to 33 °C with slices continuously perfused with standard aCSF bubbled with 95% O2 and 5% CO2. Cells were identified using diffierential interference contrast microscope (SliceScope, Scientifica, UK) with an immersion 60X objective (LUMPIanFI, Olympus, Japan) and a Photometrics IRIS9 camera (Teledyne, CA, USA). Pyramidal cells populating layer 2/3 and 5 of the auditory cortex were visually identified by their shape and distance from the cortical surface. Whole-cell currents were recorded with a Multiclamp 700B amplifier (Axon Instuments, CA, USA), low-pass filtered at 10 kHz and digitized at 20 kHz using a Digidata 1550 and pClamp software (Axon Instuments, CA, USA). Patch pipettes (open tip resistance of 3 to 5 MΩ) were pulled from glass capillaries (GC150T-10, Harvard apparatus, MA, USA), using a two-stages puller (DMZ-ZETIZ-Puller, Zeitz, Germany), and filled with an internal solution containing (in mM): 140 K-gluconate, 10 KCl, 10 HEPES, 10 Na-phosphocreatine, 4 ATP-Mg and 0.4 GTP (pH: 7.3 titrated with KOH; osmolarity 292 mOsm). All chemicals were obtained from Sigma-Aldrich (MI, USA) and ThermoFisher Scientific (MA, USA).

For the recording of light-evoked postsynaptic currents, optogenetic stimulation of OFC axons expressing ChR2 was applied on the entire field of view using a 470 nm LED (CoolLed Ltd, UK) at 5mW. Postsynaptic currents in response to 1 ms light pulse were recorded in cells clamped at -70 mV with series resistance routinely compensated. Displayed traces were obtained averaging a minimum of 5 consecutive light-evoked synaptic events after offline filtering applying a 2 kHz Gaussian low-pass filter.

### Quantification and Statistical Analysis

All analysis was done offi-line using custom written MATLAB scripts (version R2020b, MathWorks, MA,

USA).

### Behavioral performance

Behavioral performance was assessed as the percentage of correct responses calculated as ((hits + correct rejects) / total number of trials) * 100. Learning curves were obtained by fitting a psychometric curve on the behavioral performance over successive session for each mouse. Perceptual curves were obtained by fitting a psychometric curve on performance for diffierent levels of difficulty of Go/No-go comparison for each mouse. Psychometric curves were fit with a cumulative Gaussian using the open-source toolbox PsigniFit4 for Matlab ^52^. From individual learning curves we extracted the number of sessions to reach 80% of correct responses and eta, which is the overall dispersion of data points to the fitted function, as a measure of inter-session variability of performance.

### Trial-to-trial neuronal activity

Sound evoked response analysis was restricted to the time window of 0 to 250 ms from sound onset for all analysis to avoid biases related to behavioral responses that occurred later (Figure S1F).

#### Trial-to-trial variability

Sound-evoked spike counts were detected for each trial and each neuron independently for Go and No-go sounds. This resulted in a vector of sound-evoked responses for Go and No-go sounds for each neuron and for each training session. The trial-to-trial variability was characterized as the coefficient of variation of these vectors of sound evoked response for each sound type.

*Noise correlation.* Noise correlation, also called spike count correlation, was obtained by first z-scoring the vector of trial-to-trial spike counts evoked by the same stimulus: either Go or No-go sound (Figure 1) or OFC axon stimulation (Figure 6). The pairwise Pearson correlation coefficient was then used to quantify the correlation of response to the same stimulus between neurons recorded concomitantly as the measure of noise correlation.

#### Sound discriminability

The capacity of neurons to discriminate between Go and No-go sounds was assessed by computing for each training session and each neuron the Euclidean distance between z-scored vectors of trial-to-trial spike counts for Go and No-go sounds. The Euclidean distance was obtained using the *pdist2* function in Matlab. As a correlate, we computed Cosine distance using the *pdist2* function and Pearson correlation in Matlab.

### Change of selectivity across learning stages

The Selectivity Index (SI) was used to characterize the selectivity of individual neurons for either the Go or the No-go sound. The SI of an individual neuron corresponds to the diffierence between the mean response to the Go sound and the No-go sound divided by the pooled standard error ^8,28^:

A bootstrap procedure was used to detect neurons with significant change of SI with learning: for each cluster, randomly selected trials with replacement (1000 iterations) from the early stage were used to calculate corresponding SIs to create a confidence interval (2.5%-97.5%). SI at expert stage outside of this interval were considered as significantly diffierent compared to early stage.

### Optotagging

To identify SST+ neurons, blue light pulses (473nm, 10mW, 50/100ms) were presented 100 times at 1 Hz through the optic fiber cannula implanted above A1 next to the recording electrode in SST-ChR2 mice. For each cluster and each trial, the neuronal activity was aligned to the light onset. Light-responsive neurons were detected as neurons showing a significant increase of activity during light presentation compared to baseline (Student t test, *P* < 0.1), a short latency (< 10 ms, average over all trials) and a small jitter (< 5 ms) of the first spike. These neurons were considered as putative SST+ neurons.

### Dual optogenetic circuit mapping

To detect neurons responsive to OFC axon stimulation in A1, analysis was restricted to trials with only blue light. Neurons were considered responsive to the activation of OFC axons if the average evoked response to blue light illumination was higher than the baseline (mean + 2.5 SD) within 50 ms from light onset.

A bootstrap procedure was used to detect neurons with significant change of blue light-evoked response when silencing SST+ neurons (i.e. yellow light illumination): for each cluster, randomly selected trials with replacement (1000 iterations) from trials where only the blue light was presented were used to calculate corresponding mean evoked responses to create a confidence interval (10%-90%). Mean evoked response in the presence of yellow light (optogenetic SST+ neurons silencing) outside of this interval were considered as significantly diffierent.

### Statistics

Statistical analysis was conducted using MATLAB R2020b (MathWorks, MA, USA) or GraphPad Prism 9 (GraphPad Software, MA, USA). No statistical methods were used to predetermine sample size. Our sample sizes are similar to those reported in previous publications ^19,20,23^. We used non-parametric tests when residuals were not normally distributed. Statistical comparison between two groups were done using parametric two-tailed unpaired t-test (Figure 5H, 6M and 6N) or non-parametric Mann-Whitney test (Figure 4F, 5C and 6G). Comparison between two groups with paired data were done using parametric paired t-test (Figure 4E and 6I) or non-parametric Wilcoxon test (Figure 6E, F). Comparisons between more than two groups were conducted using non-parametric Kruskal-Wallis test followed by a Dunn’s post hoc multiple comparison test (Figure 5D, I, J). Comparisons between more than two groups with paired data were conducted using parametric RM-1-way ANOVA followed by a Tukey post hoc multiple comparison test (Figure 1F) or using a non-parametric Friedman test followed by a Dunn’s post hoc multiple comparison test (Figure 2F, H, J). Comparison of distributions were conducted using ꭓ^2^ test (Figure 4B, C, D). Bootstrap procedures were done as mentioned in the corresponding sections. Diffierences were considered significant at: \**P* < 0.05, \*\**P* < 0.01 and \*\*\**P* < 0.001. Data collection and analysis were not performed blind to the conditions of the experiment, but analysis relied on code that was standardized for all experimental conditions. All mice were included in the analysis unless they did not learn the task or post-hoc histological processing did not confirm proper viral injection or electrode placement.

