## Supplementary Figures for "Top-down inputs are controlled by somatostatin-expressing interneurons during associative learning"

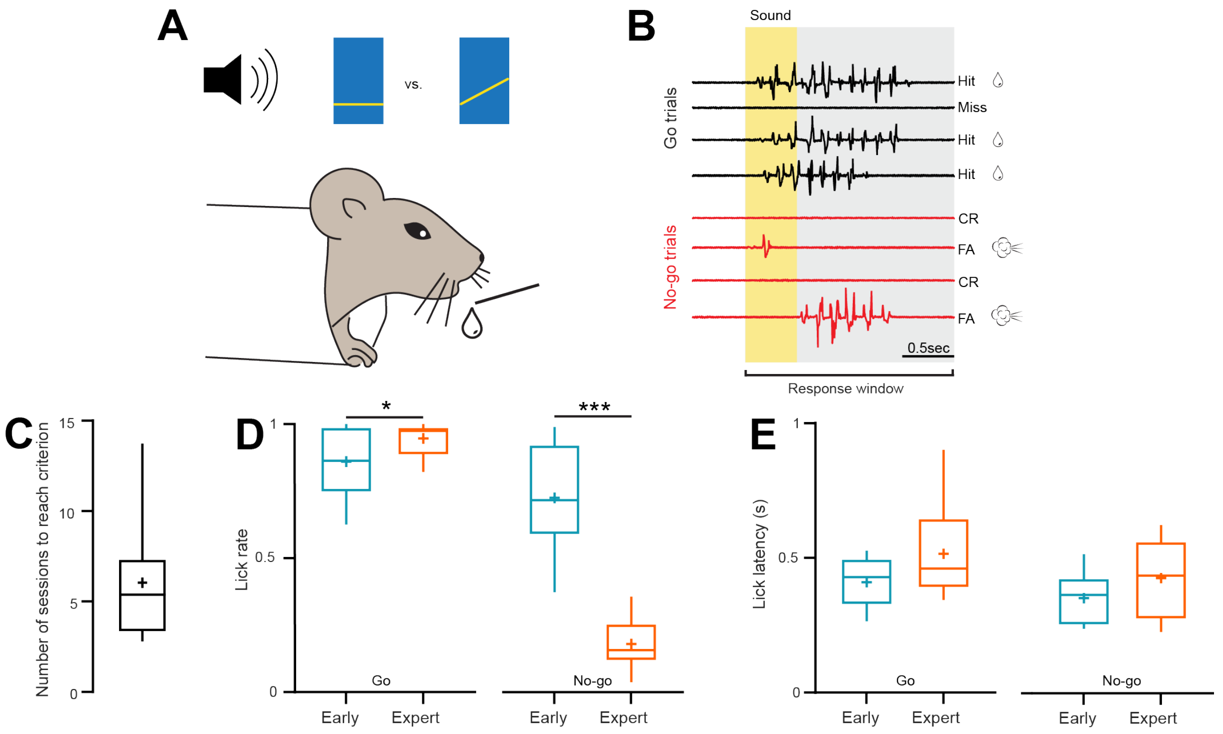


**Figure S1. Go/No-go auditory discrimination task, related to Figure 1. (A)** Schematic of a head-fixed mouse learning a Go/No-go auditory discrimination task. **(B)** Example lick traces. Sounds last 0.5 s and the response window lasts 2 s from the sound onset. Licking for the Go sound is a Hit and the mouse receives a reward. Not licking for the Go sound is a Miss. Not licking for the No-go sound is a Correct reject. Licking for the No-go sound is a False alarm and the mouse receives a punishment (mild air puff and 3 s extended time-out). **(C)** Average number of sessions to reached 80% of correct responses. **(D)** Comparison of the lick rate in response to the Go sound (left) and No-go sound (right) at early and expert stage. (Go: $P$=0.0174, No-go: $P$<0.0001). **(E)** Comparison of the lick latency in response to the Go sound (left) and the No-go sound (right) at early and expert stage. (n=10, paired t-test, *p < 0.05, ***p < 0.001). Boxplots represent the min, 25$^{th}$ percentile, median, 75$^{th}$ percentile, and max; + represent the means.

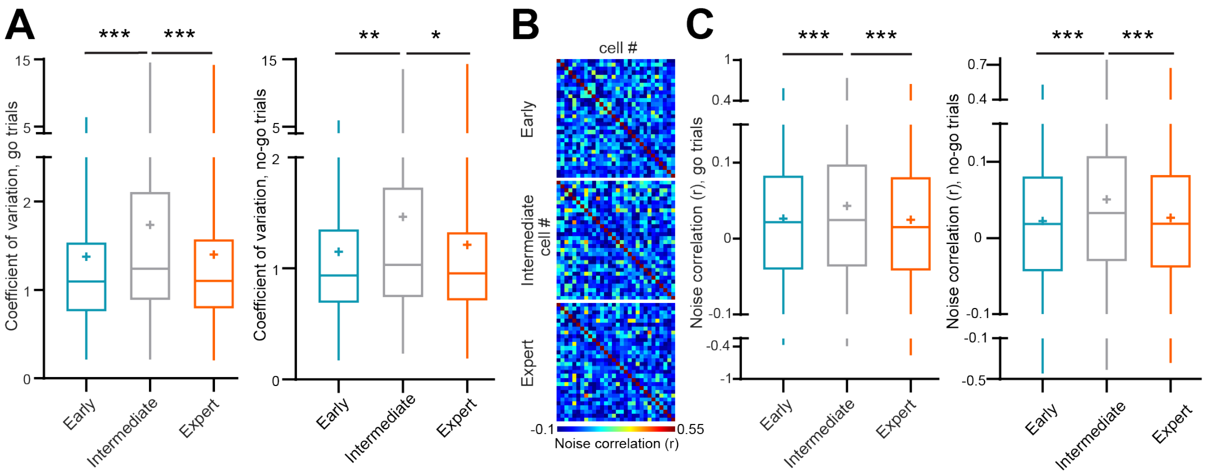


**Figure S2. Associative learning increases trial-to-trial variability of sound evoked responses, related to Figure 1. (A)** Coefficient of variation of trial-to-trial sound evoked responses (0-250 ms from sound onset) at three stages of learning. (Left: Go tones, $F(2, 409)=8.925$, $P=0.0002$; post hocs: early vs. intermediate: $P=0.0008$, intermediate vs. expert: $P=0.0009$; Right: No-go tones, $F(2, 410)=6.991$, $P=0.001$; post hocs: early vs. intermediate: $P=0.0012$, intermediate vs. expert: $P=0.0153$). **(B)** Example correlation matrix from the same mouse over learning stages for the Go tone. **(C)** Noise correlation as a function of learning stages (Left: Go tones, $F(2, 8992)=42.37$, $P<0.0001$; post hocs: early vs. intermediate: $P<0.0001$, intermediate vs. expert: $P<0.0001$; Right: No-go tones, $F(2, 9059)=102.4$, $P<0.0001$; post hocs: early vs. intermediate: $P<0.0001$, intermediate vs. expert: $P<0.0001$). (n=5 mice, 207 neurons, RM one-way ANOVA with Tukey's multiple comparison test, * $p < 0.05$, ** $p < 0.01$, *** $p < 0.001$). Boxplots represent the min, 25$^{th}$ percentile, median, 75$^{th}$ percentile, and max; + represent the means.

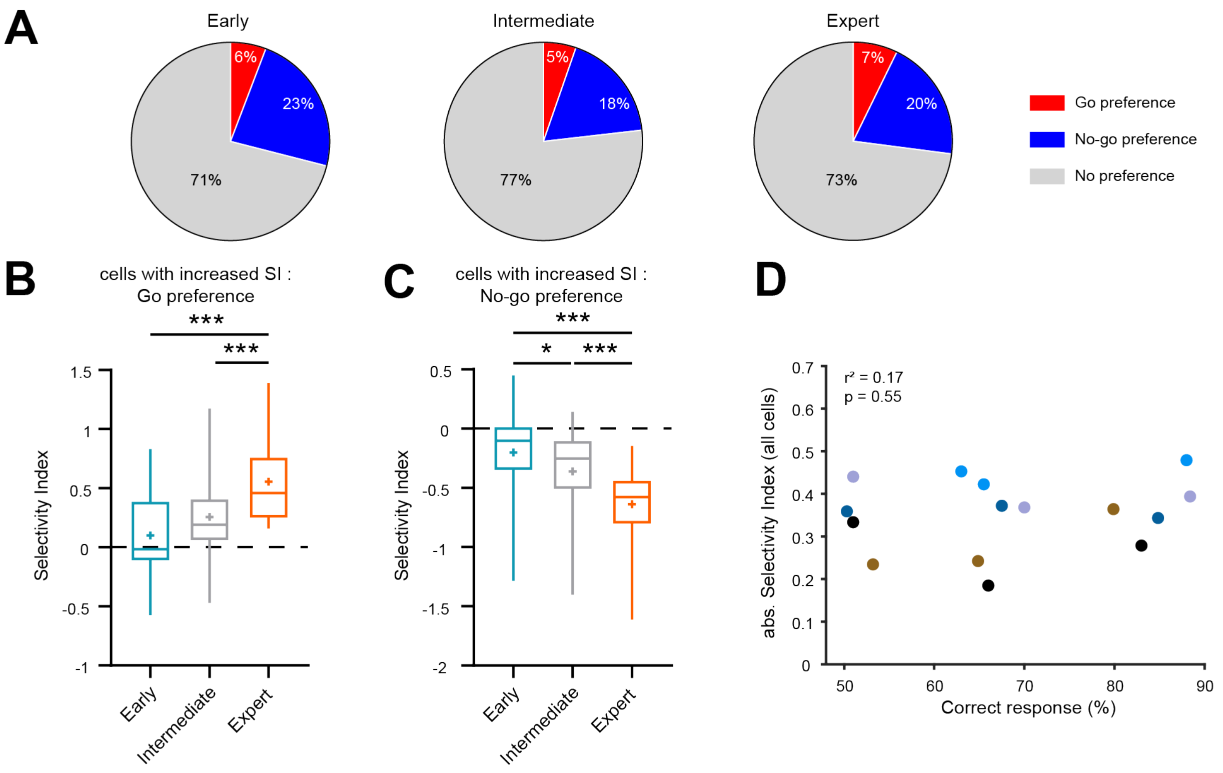


**Figure S3. Associative learning increases the selectivity of a subset of neurons to the learned cues, related to Figure 2. (A)** Distribution of neurons based on their preferred selectivity to Go or No-go sound. **(B)** SI of cells with a significant change of selectivity over learning that are selective for the Go sound at expert stage. (*P*<0.0001, post hocs: early vs. expert: *P*<0.0001, intermediate vs. expert: *P*=0.0009). **(C)** SI of cells with a significant change of selectivity over learning that are selective for the No-go sound at expert stage. (*P*<0.0001, post hocs: early vs. intermediate: *P*=0.0276, early vs. expert: *P*<0.0001, intermediate vs. expert: *P*<0.0001). **(D)** Correlation between the mean absolute selectivity index of neurons for each mice and the behavioral performance for each mice and each learning stage (Pearson correlation). Each color represents an individual mouse. (n=5 mice, 207 neurons, Friedman test, * p < 0.05, *** p < 0.001). Boxplots represent the min, 25th percentile, median, 75th percentile, and max; + represent the means.

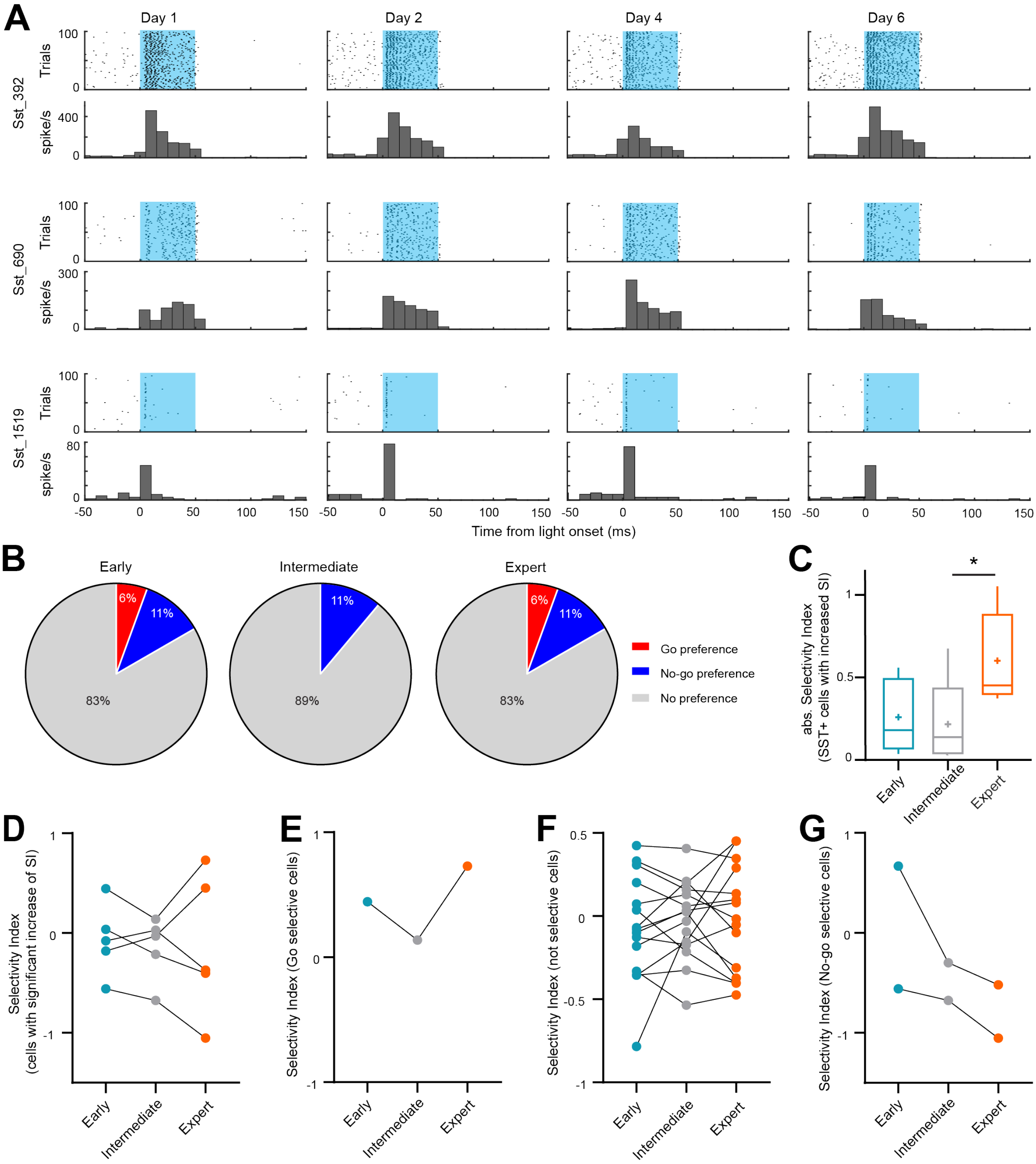


**Figure S4. Selectivity index of SST+ neurons across learning, related to Figure 3. (A)** Three example light-responsive neurons (SST expressing ChR2) recorded over multiple days. Blue shaded areas represent blue light presentation (473 nm, 10 mW, 50 ms). **(B)** Distribution of neurons based on their preferred selectivity to Go or No-go sound. **(C)** Mean absolute SI of neurons with a significant increased selectivity during learning. (Friedman test: *P*=0.0239, Dunn's multiple comparison test: *P*=0.0342). **(D)** Selectivity index of SST+ neurons with a significant increase of selectivity over learning. **(E)** Selectivity index of SST+ neurons selective for Go sound (SI > 0.5) at expert stage as a function of learning stages. **(F)** Same as e for neurons with no preference (-0.5 < SI > 0.5) at expert stage. **(G)** Same as E for neurons with No-go preference (SI < -0.5) at expert stage. (n=4 mice, 18 neurons).

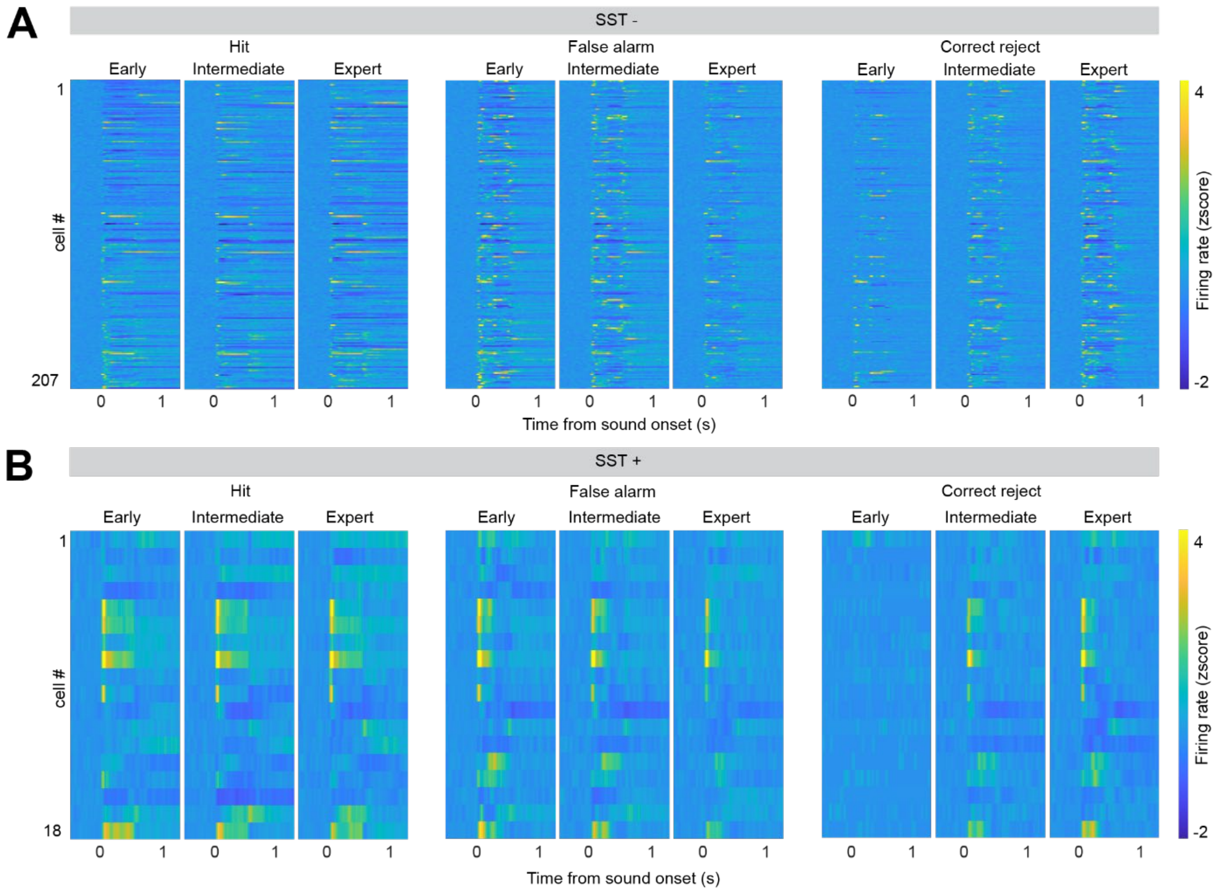


**Figure S5. Neuronal response to sounds split based on task-outcome categories, related to Figure 4.** **(A)** Heatmaps of SST– neurons for Hit trials (left), False alarm trials (middle) and Correct reject trials (Right). Neurons are sorted as in Figure 4. **(B)** Same as A. for SST+ neurons.

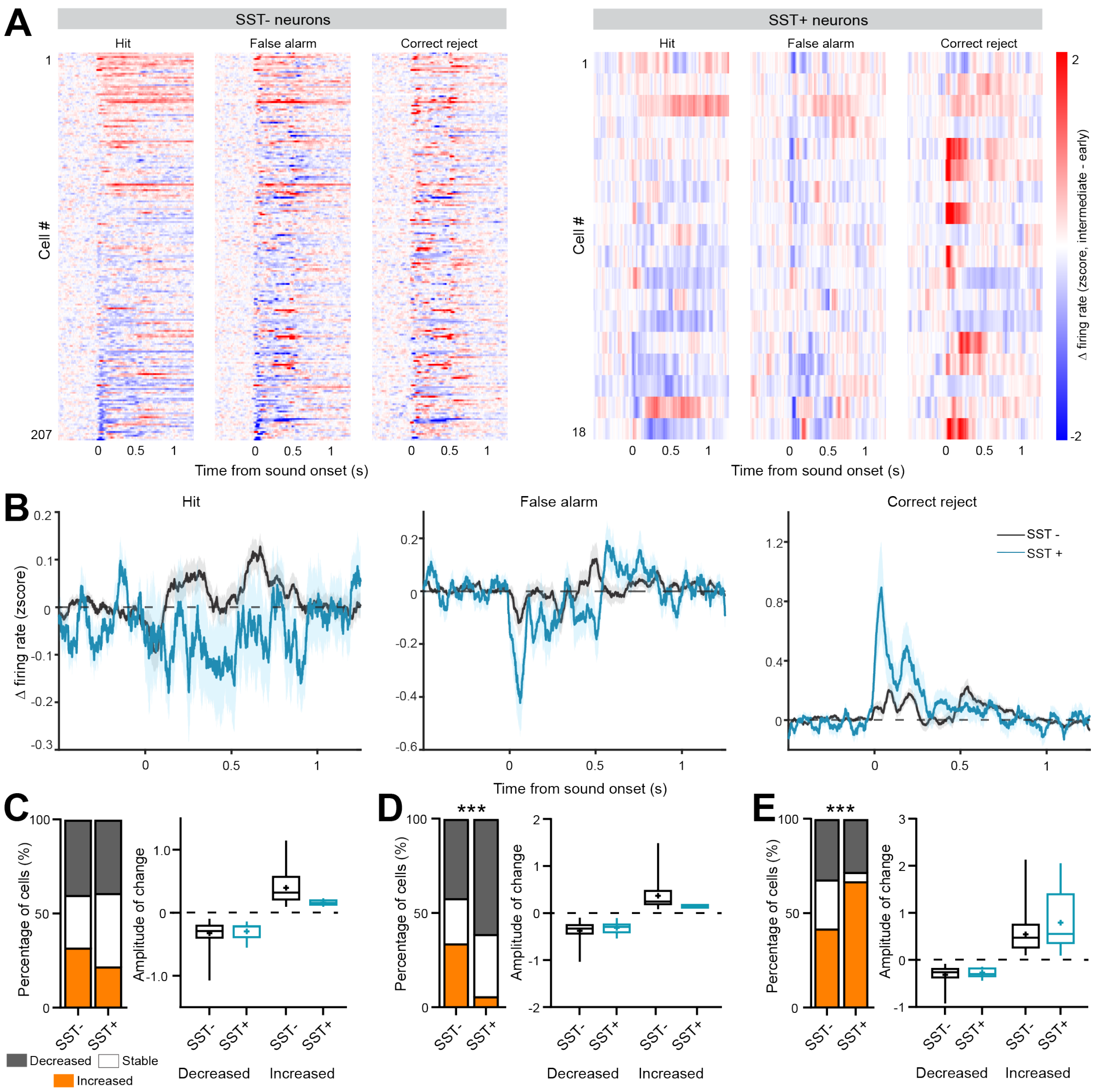


**Figure S6. Learning-induced neuronal plasticity from early to intermediate stage is characterized by opposite changes of SST+ and SST- neurons, related to Figure 4. (A)** Difference of peristimulus time histogram between the early stage and the intermediate stage for each neuron segregated between Hit trials, False alarm trials and Correct reject trials for SST- neurons (left, 207 neurons) and SST+ neurons (right, 18 neurons). Heatmaps are sorted based on maximum change of response in Hit trials. **(B)** Average ΔPSTH for Hit trials (left), False alarm trials (middle) and Correct reject trials (right). **(C)** Percentage of neurons with changed response (left) and amplitude of response's change (right) for Hit trials. **(D)** Same as C for False alarm trials. (*P*<0.0001). **(E)** Same as C for Correct reject trials. (*P*<0.0001). (SST- n=207, SST+ n=18, $\chi^2$, *** p < 0.001).

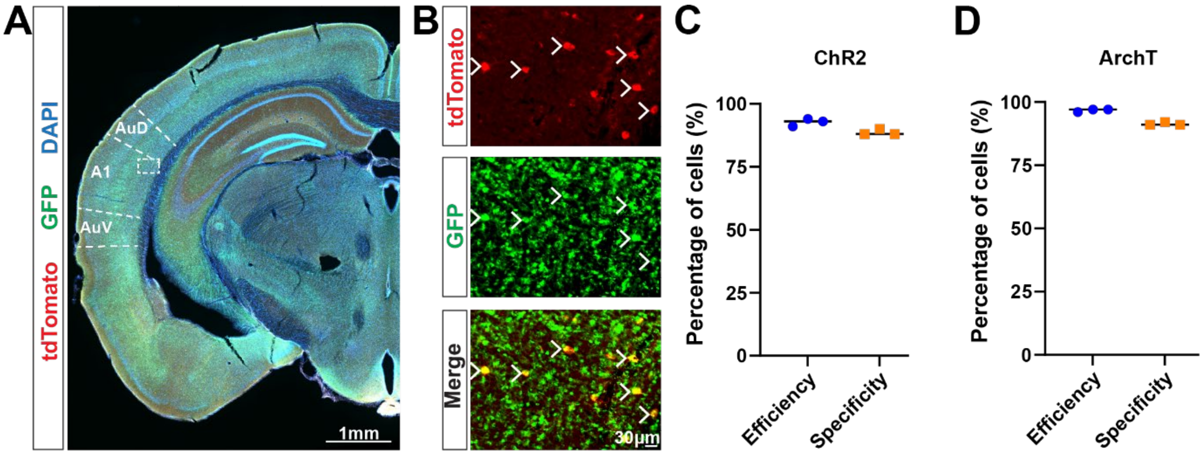


**Figure S7. Characterization of the expression of opsins in SST+ neurons, related to Figure 5. (A)** Exemple brain slice stained for tdTomato (SST), green fluorescent protein (GFP, opsins) and DAPI (cell nucleus). **(B)** Inset from (a) showing an enlargement of the co-expression of tdTomato and GFP (arrows) in SST+ neurons. **(C)** Efficiency (number of cells SST+ that express opsins: (tdTomato+ & GFP+)/ tdTomato+) and specificity (number of cells expressing opsins that are SST+: (tdTomato+ & GFP+)/ GFP+) of the expression of ChR2 in SST+ neurons in SST-ChR2 mice. **(D)** Efficiency and specificity of the expression of ArchT in SST+ neurons in SST-ArchT mice.

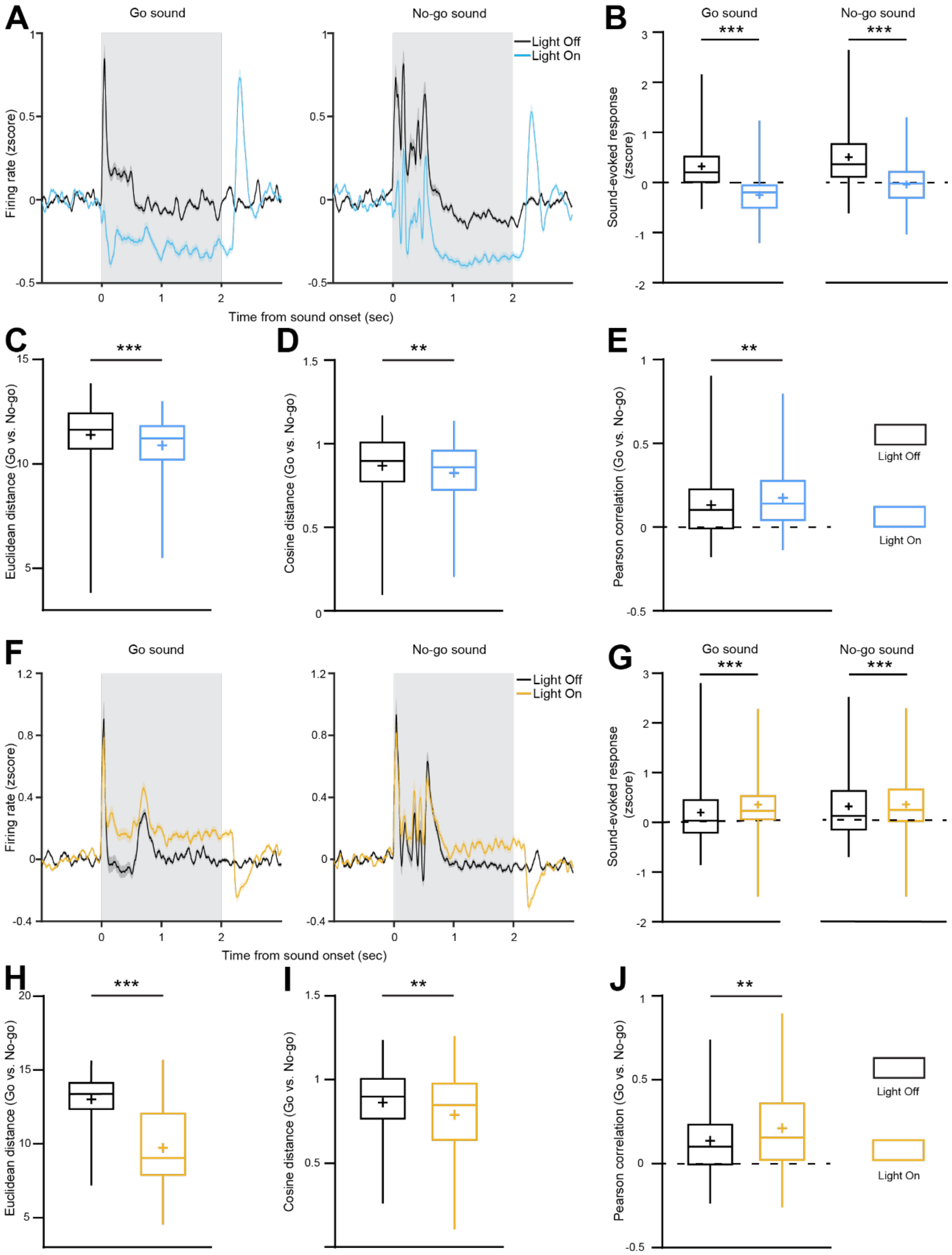


**Figure S8. Effect of photomodulation of SST+ neurons in A1 on sound processing, related to Figure 5.** **(A)** Average PSTH for Go sound (left) and No-go sound (right) with blue-light activation of SST+ neurons in SST-ChR2 mice (blue, n=207) or without light (black). **(B)** Mean sound-evoked response for the Go and No-go sound with optogenetic activation of SST+ neurons (blue) or without (black). (Go: *P*<0.0001, No-go: *P*<0.0001). **(C)** Effect of SST+ neurons optogenetic activation on the mean Euclidean distance between trial-to-trials sound evoked response for the Go or the No-go sound. (*P*<0.0001). **(D)** Effect of SST+ neurons optogenetic activation on the mean cosine distance between trial-to-trials sound evoked response for the Go or the No-go sound. (*P*=0.0067). **(E)** Effect of SST+ neurons optogenetic activation on the mean Pearson correlation between trial-to-trials sound evoked response for the Go or the No-go sound. (*P*=0.0067). **(F-J)** Same as A-E for SST+ silencing in SST-ArchT mice (n=190). (g, Go: *P*<0.0001, No-go: *P*<0.0076. h, *P*<0.0001. i, *P*=0.0016. j, *P*=0.0016. Mann-Whitney test). Boxplots represent the min, 25$^{th}$ percentile, median, 75$^{th}$ percentile, and max; + represent the means.

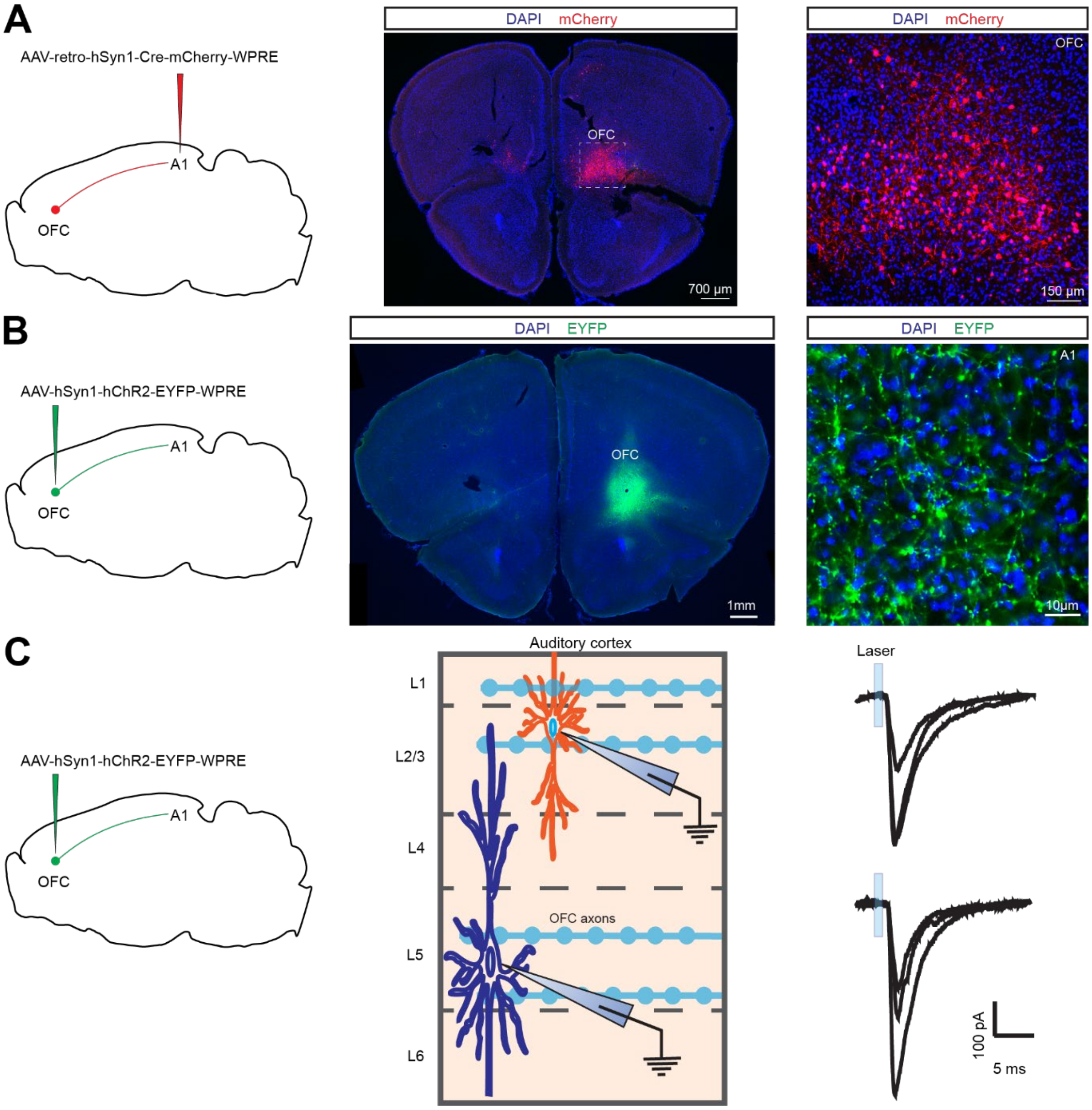


**Figure S9. Identification of direct projections from OFC to A1, related to Figure 6. (A)** Schematic of the injection of a retrograde AAV in A1 to visualize A1-projecting neurons in OFC (left). Brain slice with the OFC showing A1-projecting neurons in red (middle) with a magnification (right) corresponding to the inset. **(B)** Schematic of the injection of an AAV in OFC to identify OFC neurons' axons in A1 (left). Brain slice showing the injection site in OFC (middle). Axons of OFC neurons in A1 (right). **(C)** Schematic of slice electrophysiology recording of the response of A1 neurons to the stimulation of OFC neurons' axons expressing ChR2 (left and middle). Average post-synaptic blue light-evoked currents recorded in L2/3 pyramidal neurons (right top) and L5/6 pyramidal neurons (right bottom) (10-15 repetitions, n=3 neurons each).

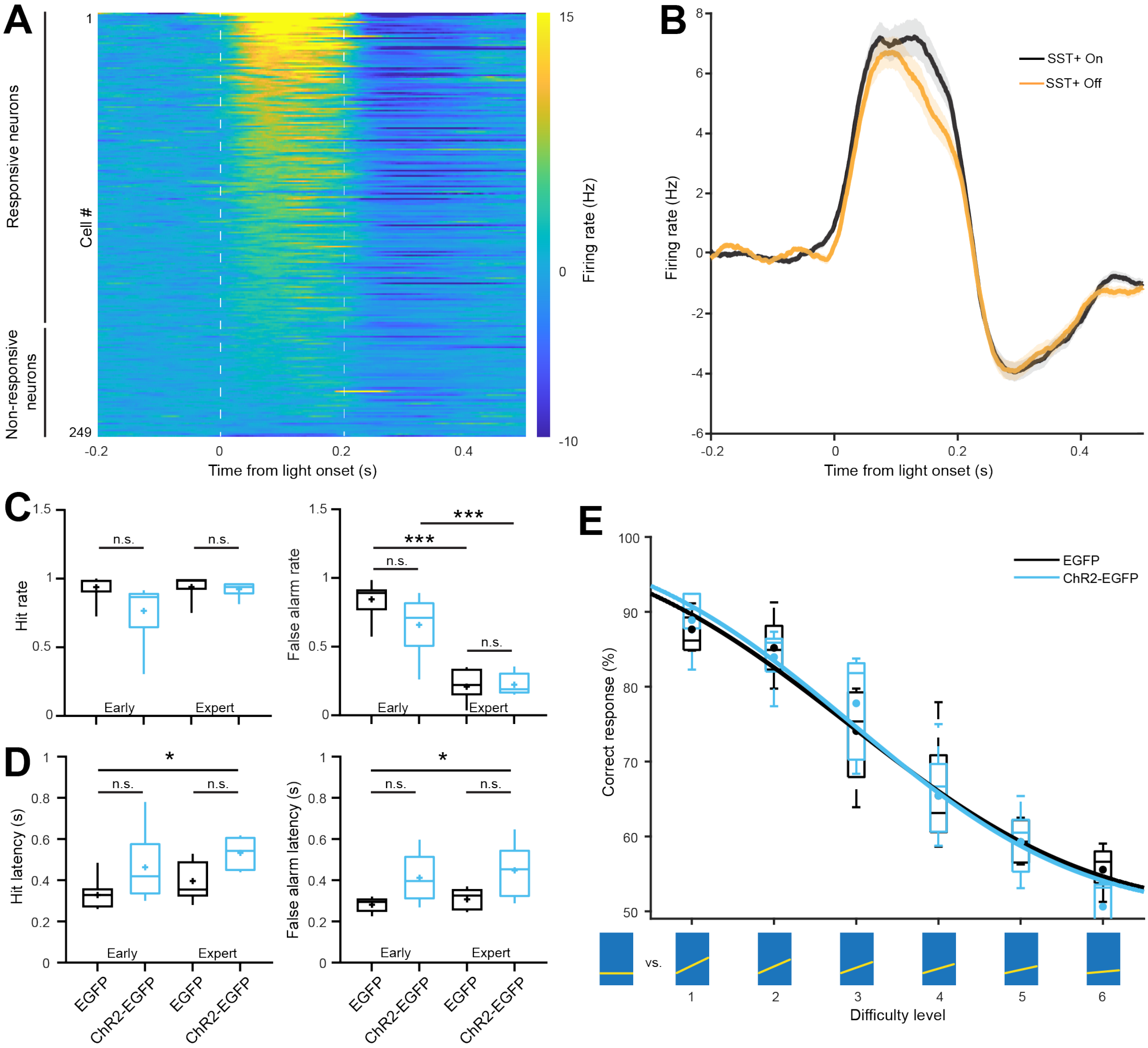


**Figure S10. Modulation of OFC inputs to A1 by SST+ neurons, related to Figure 6. (A)** Heatmap of the response of A1 neurons to the optogenetic stimulation of OFC axons. **(B)** Mean response of A1 blue-light responsive neurons to the optogenetic stimulation of OFC axons during optogenetic silencing of SST+ neurons (yellow) or no modulation of SST+ neurons (black) (n=174). **(C)** Comparison of the response rate for Hit trials and False alarm trials in mice expressing ChR2 in OFC axons (blue, n=6) and control mice (black, n=7) at early and expert stage. (*P*<0.0001 One-way ANOVA test, *P*<0.0001 and *P*=0.0002 Tukey multiple comparison test). **(D)** Comparison of the response latency for Hit trials (*P*=0.0228 One-way ANOVA test, *P*=0.0181 Tukey multiple comparison test) and False alarm trials (*P*=0.0119 One-way ANOVA test, *P*=0.0218 Tukey multiple comparison test) in mice expressing ChR2 in OFC axons (blue, n=6) and control mice (black, n=7) at early and expert stage. Boxplots represent the min, 25$^{th}$ percentile, median, 75$^{th}$ percentile, and max; + represent the means. **(E)** Psychometric function of the performance in the perceptual task for mice expressing ChR2 in OFC axons (blue, n=6) and control mice (black, n=7). Boxplots represent the 25$^{th}$ percentile, median, 75$^{th}$ percentile, and ±s.e.m; + represent the means.
